# Sex-specific lymphoid lineage differentiation is modulated by CIZ1

**DOI:** 10.64898/2026.08.26.747233

**Authors:** Elena Guglielmi, Ramiro Monge-Lozano, Joanna F. Pearson, Fabiano S. Pais, Sally R. James, James P. Hewitson, David G. Kent, Dawn Coverley, Justin F.X. Ainscough

## Abstract

The epigenetic stability factor CIZ1 helps maintain X-chromosome inactivation, and its absence results in murine female-specific splenomegaly. By exploring the aetiology of this pathology, we reveal unexpected evidence for dosage compensation via modification of X-linked gene expression in males. Genes known to escape repression on the inactive X-chromosome in females, including the B-cell maturation factor DDX3X, require CIZ1 in males to maintain parity between the sexes. Furthermore, absence of this single regulator triggers stark sex-specific changes to large autosomal domains in B cells, causing naive female B cells to shift prematurely towards germinal centre transcriptional signatures, including immunoglobulin and pro-proliferation genes even without immune challenge. Conversely, males acquire natural killer-like gene expression through elevation of Killer cell Lectin-like Receptors. Together, the data indicate that CIZ1 limits sexually dimorphic gene expression on autosomes, and promotes dosage compensation by modulation of the male X-chromosome, introducing a new paradigm for sex biased disorders of the immune system.

**Highlights:**

- CIZ1 forms protein assemblies at the inactive X-chromosome in females, that disaggregate in naive B cells and reform on antigen stimulation *in vitro* and challenge *in vivo*.
- X-linked escape gene expression in females is normally compensated in males, but polarises in the absence of CIZ1 via suppression in males.
- Autosomal gene clusters respond to absence of CIZ1 in a sex-specific manner leading to divergent B cell differentiation even before immune challenge.

**Summary:** This study shows that CIZ1 promotes dosage compensation of X-linked escape genes via modulation of male gene expression, and also dampens divergent expression of autosomal loci in B cells, protecting against sex-biased gene expression, and lymphocyte hyperplasia in females.

**Graphical abstract:** 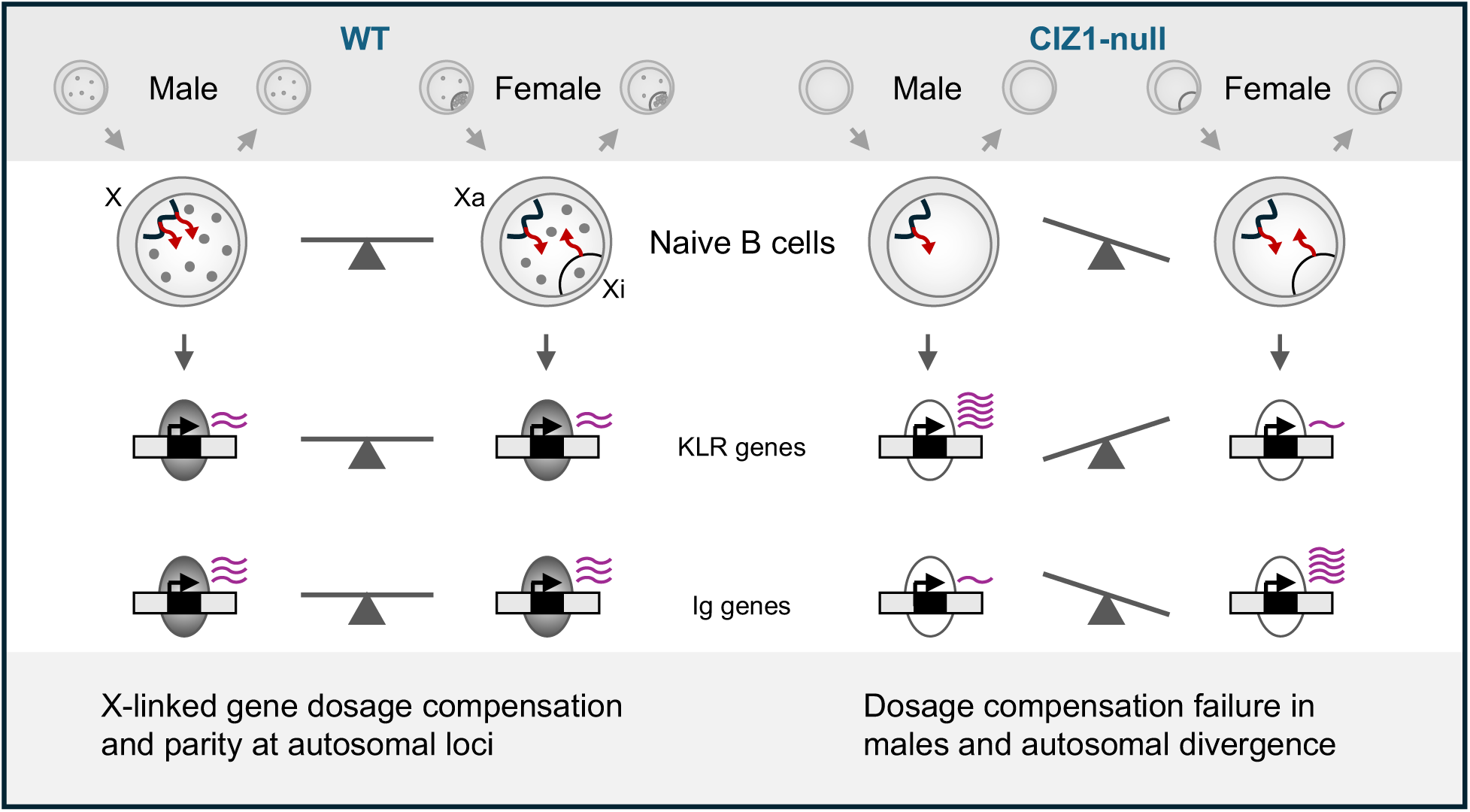

## INTRODUCTION

Sex is a biological variable that influences the immune system, affecting both innate and adaptive immune responses, resulting in differential susceptibility of males and females to infectious and autoimmune diseases, organ rejection rates, and cancer (Dunn et al., 2024; Forsyth et al., 2024; Libert et al., 2010). The sex chromosomes (XX in females and XY in males) are thought to underlie sex-biased immune cell function through multiple mechanisms, including elevated expression of X-linked immune genes (Radovanovic et al., 2026), and the suggestion that *Xist* ribonucleoprotein (RNP) complexes may act as autoantigens (Dou et al., 2024).

Transcriptional repression of one copy of the X-chromosome in female mammals equalises the expression of most X-linked gene products between females and males (Loda et al., 2022; Lyon, 1961). The long non-coding RNA (lncRNA) *Xist* initiates X-chromosome inactivation (XCI) by nucleating large supramolecular assemblies that recruit enzymes that modify DNA and histones to heritably compress chromatin and repress transcription from the inactive X-chromosome (Xi) (Brockdorff et al., 1992; Brown et al., 1992; Clemson et al., 1996; Markaki et al., 2021; Penny et al., 1996).

Once established in the embryo, most female somatic cells maintain XCI during differentiation, however not all Xi genes are stably repressed as some continue to require expression of *Xist* and the ongoing recruitment of repressive chromatin modifiers (Loda et al., 2022; Wutz and Jaenisch, 2000). In mice 3–7% of genes ‘escape’ silencing in most cell types while a further ∼20% escape silencing in some cell types (Hauth et al., 2026; Peeters et al., 2023; Yang et al., 2010), leading to lack of full dosage compensation and potentially sexually dimorphic phenotypes.

In haematopoietic stem and progenitor cells and activated B and T lymphocytes *Xist* is enriched around Xi chromatin. However, in mature naive lymphocytes from humans and mice, *Xist* assemblies and some repressive histone post-translational modifications (PTMs) are lost from Xi, despite continued production of *Xist* during this window. Nevertheless, for the majority of genes, repression that was established earlier in development persists (Grigoryan et al., 2021; Ridings-Figueroa et al., 2017; Savarese et al., 2006; Syrett et al., 2019; Wang et al., 2016). The biological significance of this unusual behaviour of *Xist* in the haematopoietic lineage is not understood, and also not straightforward to study *in vivo*. Evidence *in vitro* shows that antigen stimulation of naive lymphocytes leads to the rapid accumulation of *Xist* particles and repressive histone post-translational modifications (PTMs) in the Xi territory, which is thought to reflect the *in vivo* response to immune challenge (Ridings-Figueroa et al., 2017; Syrett et al., 2019; Wang et al., 2016).

Cip1-Interacting Zinc Finger protein 1 (CIZ1) interacts directly with *Xist* and accumulates in large RNA-protein assemblies around Xi chromatin (Dixon-McDougall and Brown, 2022; Markaki et al., 2021; Ridings-Figueroa et al., 2017; Rodermund et al., 2021; Sofi et al., 2022; Sunwoo et al., 2017; Valledor et al., 2023). In fibroblasts, its absence is sufficient to cause dispersal of *Xist*, depletion of repressive histone PTMs, and changes in the expression of some X-linked genes (Dobbs et al., 2023; Ridings-Figueroa et al., 2017; Stewart et al., 2019; Turvey et al., 2025). In addition, CIZ1 also forms smaller RNA-dependent assemblies throughout the nucleus in both sexes, and its absence disrupts the expression of hundreds of genes on autosomes, indicating roles beyond the X chromosome (Stewart et al., 2019; Turvey et al., 2025). CIZ1-*Xist* assemblies are dependent on two RNA-interaction domains in the N- and C-terminus of CIZ1 (Sofi and Coverley, 2023), which can be disrupted experimentally causing rapid changes in underlying chromatin and gene expression, that is mirrored in early-stage human breast cancers (Byrom et al., 2026; Turvey et al., 2025). Together, the data indicate that localized CIZ1 assemblies help create spatially constrained environments that affect chromatin state (Fig.1A).

**Fig. 1.**
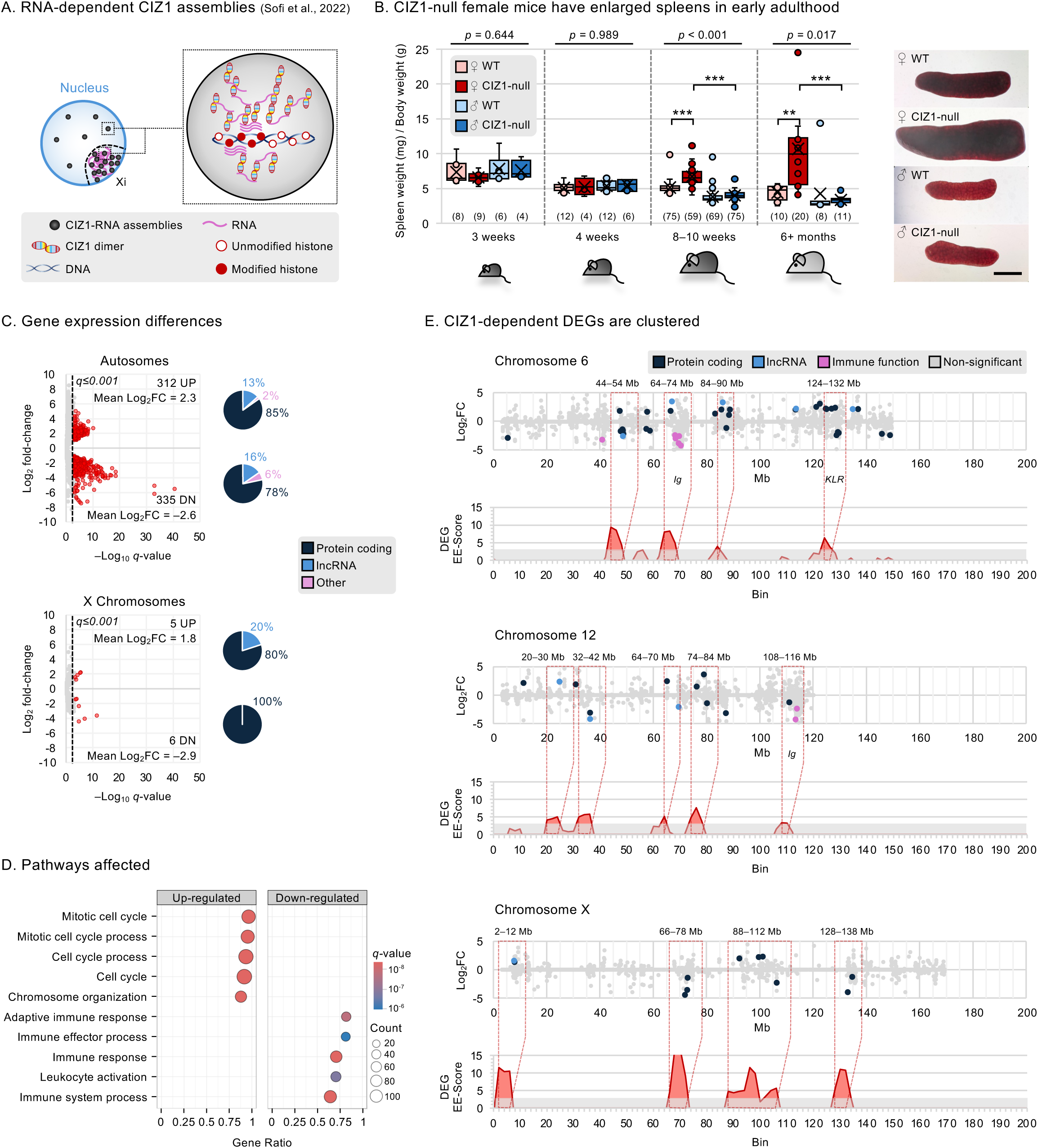
CIZ1-dependent gene clusters in female spleen A) Schematic showing RNA-dependent CIZ1 protein assemblies protecting histone post-translational modifications in underlying chromatin, evidenced by analysis of Xi (Sofi et al., 2022; Turvey et al., 2025). B) Spleen weight index showing number of animals in parentheses, and significant differences (Two-Way ANOVA). Right, example images showing spleen morphology at 6 months of age. Scale bar is 5 mm. C) Volcano plots showing gene expression in spleen tissue, comparing three WT to three CIZ1-null female mice at 15+ months of age, for autosomes and the X chromosomes. Differentially expressed genes (red, *q*≤0.001) and biotype distribution of up-regulated (UP) and down-regulated (DN) genes that are changed in CIZ1-null compared to WT are shown. D) The top five UP and DN gene expression pathways (GSEA M5 biological processes) affected by absence of CIZ1 (autosomal and X-linked differentially expressed genes combined). E) Chromosomal locations of DEGs on chromosomes 6, 12, and X, highlighting protein coding genes (dark blue), lncRNAs (light blue), immune-related genes (pink), and non-significantly affected genes (grey). Below, for each chromosome DEG clusters (red) are defined by an inclusion threshold of EE-Score≥3.

At the organismal level, absence of CIZ1 has subtle and diverse effects that show sex bias. CIZ1-null mice develop normally with no overt defects in embryogenesis or early postnatal development arguing that, unlike *Xist,* CIZ1 is not essential for establishment of XCI. However, in adulthood a lymphoproliferative disorder emerges specifically in females (Ridings-Figueroa et al., 2017; Sunwoo et al., 2017). Assessment of behavioural phenotypes in the same model revealed motor impairment that was worse in females, and cognitive impairment in both sexes (Khan et al., 2018; Xiao et al., 2016), while human studies link CIZ1 loss with female-specific neurodevelopmental defects (Besnard et al., 2025). While these data argue for sex-specific roles in multiple tissue lineages, we focus here on the immune system as a manipulable model of cellular transition. The sensitivity of the blood compartment to deletion of *Xist* (Radovanovic et al., 2026; Yang et al., 2022; Yildirim et al., 2013), and the unusual behaviour of *Xist* in lymphocytes adds additional dimensions to the question of why is the absence of CIZ1 sufficient to cause splenic hyperproliferation and why only in females.

## RESULTS

### Spleen dysplasia in adult female mice

Early analysis of phenotypic abnormalities in CIZ1-null mice generated via an insertional mutagenesis strategy (Supplemental Fig.1A), reported splenomegaly at 15+ months of age, but only in females (Ridings-Figueroa et al., 2017). Female mice presented with enlarged secondary lymphoid tissues (spleen and lymph nodes) and infiltration of abnormal B and reactive T lymphocytes similar to non-Hodgkin follicular-type lymphoma (Ridings-Figueroa et al., 2017). This striking result is corroborated here in younger mice, showing consistently enlarged spleen weight and size relative to body weight. CIZ1-null males remain largely unaffected confirming that the abnormality exhibits a strong sex-bias (Fig.1B, Supplemental Table 1).

Comparison of the transcriptomes of spleen tissue from three WT and three CIZ1-null female mice at 15+ months of age revealed 647 autosomal and 11 X-linked differentially expressed genes (DEGs, FDR q≤0.001) that are changed in CIZ1-null relative to WT. Both up-regulated (UP) and down-regulated (DN) DEGs show similar proportions of biotypes and are predominantly protein coding (Fig.1C, Supplemental Data set 1). Gene Set Enrichment Analysis of Biological Processes (GSEA GO:BP) revealed significant over representation of genes linked with cell division among UP DEGs, while the DN DEGs returned regulation of immune cell activation and proliferation (Fig.1D, Supplemental Data set 1). This correlates absence of CIZ1 with a shift towards pro-proliferation gene expression.

### CIZ1-dependent genes are spatially clustered

The 647 autosomal DEGs are not distributed evenly throughout the genome, but appear clustered, most often in regions of high gene density (Fig.1E). To quantify the non-random distribution of CIZ1-dependent genes and compensate for gene density, we compared DEG enrichment to gene enrichment. Briefly, the number of genes and the number of DEGs in 6Mb bins (slid by 2Mb) were calculated for each chromosome, and the degree of enrichment over the chromosomal average used to return individual enrichment scores for each bin. Bins were classified as DEG-enriched when their DEG enrichment score was ≥3 higher than their gene enrichment score, giving rise to a DEG Excess Enrichment Score (EE-Score). This classification identified bins on all chromosomes encompassing CIZ1-regulated genes, describing chromosomal domains that span tens of Mbs (Fig.1E, Supplemental Fig.1B,C, Supplemental Data set 1). Overall, this aligns with studies in primary human breast cancers and mouse fibroblasts, which also showed that DEGs affected by disruption of CIZ1 are clustered within large susceptible chromosomal domains (Turvey et al., 2025).

In splenocytes, some DEG-defined cluster sites encode gene families, notably the killer cell lectin-like receptor (KLR) gene cluster on chromosome 6 (∼130 Mb), and immunoglobulin gene (light and heavy chain Igv) clusters on chromosomes 6 (∼70 Mb) and 12 (∼113–115 Mb) (Fig.1E). Similar to autosomes, the X chromosome also harbours DEG-defined cluster sites (Fig.1E), which in our model could be specified by transcripts emanating from the active or inactive X.

While these data consolidate the idea of CIZ1-sensitive chromosomal domains in haematopoietic lineages, the advanced phenotype of the adult-derived spleen tissue means that they do not necessarily report on gene expression changes that drove the phenotype, and could be reporting on a historic shift in cell populations.

### B cell defects in young females

To focus investigation on earlier events in younger animals we established that splenomegaly becomes evident in CIZ1-null females, but not males, after 4 weeks and before 8-10 weeks of age (Fig.1B, Supplemental Table 1). Flow cytometry of total splenocytes isolated from 8–10 week old females revealed that CIZ1-null spleens already contain over 30% more total live cells (excluding red-blood cells) than WT, consistent with the observed spleen enlargement (Fig.2A,B). Gating on B cells (B220^+^ CD19^+^), T helper cells (T_h_; TCRb^+^ CD4^+^ CD8^−^) and T cytotoxic cells (T_c_; TCRb^+^ CD4^−^ CD8^+^) showed that increase in the B cell fraction in CIZ1-null compared to WT is largely responsible. A small but significant increase was also seen in T_h_ cells, which can help B cell responses including affinity maturation and antibody isotype class switching. Elevated B and T cells in young mice is consistent with onset of the adult phenotype of non-Hodgkin-type B cell lymphoma with reactive T cell infiltrates (Ridings-Figueroa et al. (2017).

**Fig. 2.**
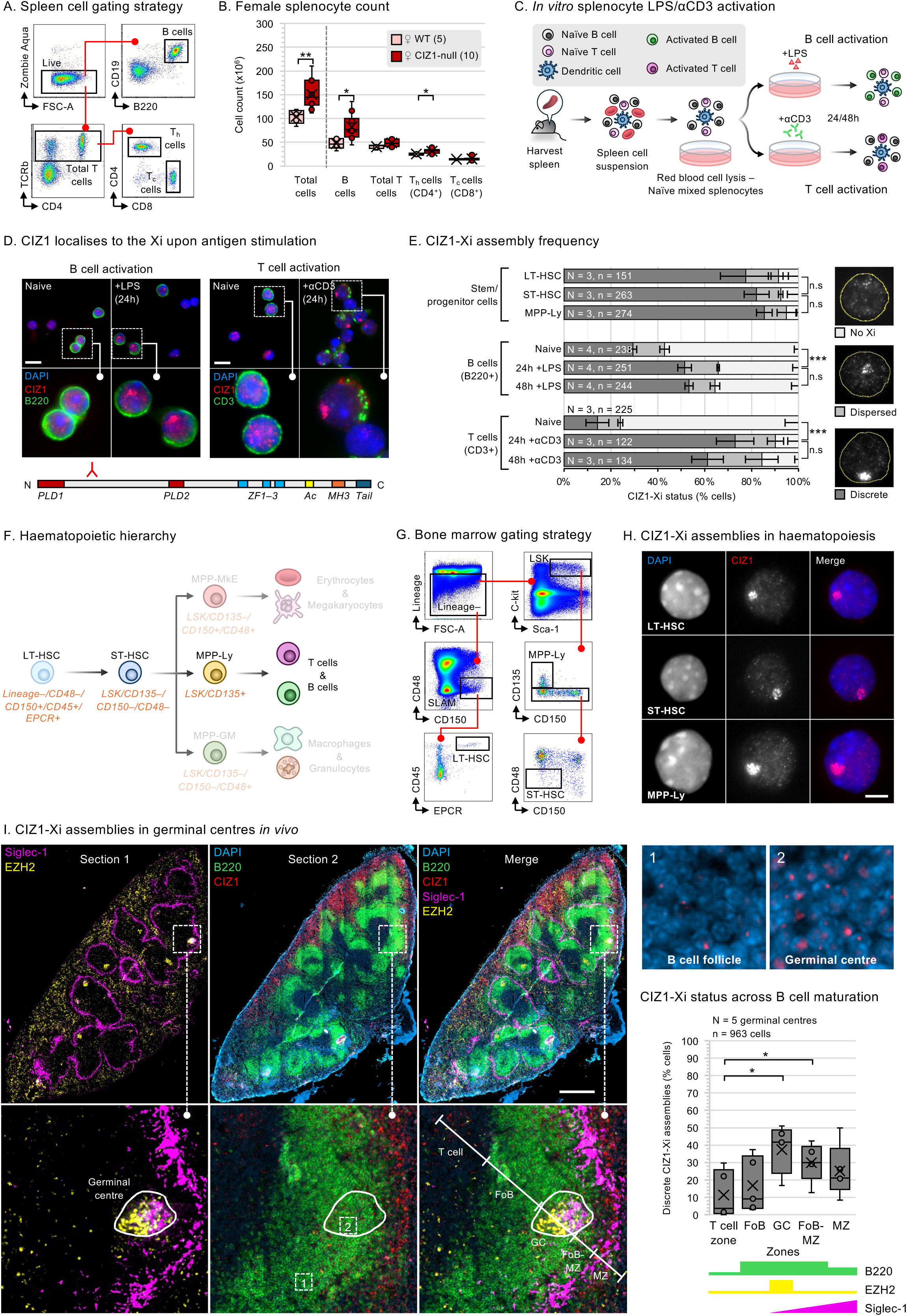
Dissipation and reformation of CIZ1-Xi assemblies defines a window in lymphocyte differentiation A) Gating strategy for sorting B and T (T_h_ and T_c_) cells from splenocytes by flow cytometry. B) Splenocyte counts from five WT and 10 CIZ1-null female mice at 8–10 weeks of age. For total cells *p*=0.00814, B cells *p*=0.0185, and CD4 T_h_ cells *p*=0.036, Student’s T-test. C) *In vitro* B and T cell activation strategy, using LPS or αCD3 for 24 or 48 hours. D) (Top) Immuno-fluorescence images on 8–10 week old female WT splenocytes showing CIZ1 (red) and B220^+^ B cells or CD3^+^ T cells (green), and DAPI (blue). Scale bar is 10 µm. Insets depict lack of and re-aggregation of CIZ1 assemblies upon activation. (Bottom) Schematic of CIZ1 protein domains (Byrom et al., 2026), showing region recognised by anti-CIZ1 rabbit polyclonal antibody 1794. E) Quantification of CIZ1-Xi assembly status in the indicated cell types, as fully formed (discrete), partially formed (dispersed) or absent (no Xi assembly), as illustrated on the right. N, number of independent cell isolates (mice) and n, number of cells. Significance indicators from Student’s T-test performed on discrete CIZ1-Xi populations. F) Haematopoietic cell hierarchy with focus on the lymphoid lineage (centre). Cell surface markers used to discriminate cell types are shown. LSK refers to Lineage^−^ Sca-1^+^ c-kit^+^. G) Strategy for gating LT-HSC, ST-HSC, and MPP-Ly from bone marrow by flow cytometry. H) Example images of sorted LT-HSC, ST-HSC, and MPP-Ly cells, showing CIZ1 (red) and DAPI (blue). Scale bar is 5 µm. I) Whole spleen 5 µm transverse cryosections from a 9-week-old WT female mouse. Section 1 was stained for Siglec-1 (purple), and EZH2 (yellow), and adjacent section 2 for DAPI (blue), B220 (green) and CIZ1 (red). Images were merged to create a five-colour analysis. Scale bar is 500 µm. Numbered insets in the lower middle panel show boxed regions 1 (B cell follicle) and 2 (germinal centre), enlarged on the right showing CIZ1 (red) and DAPI (blue). Below, quantification of CIZ1-Xi status of cell populations contacting the line traversing the indicated zones: T cell zone, follicular B cell zone (FoB), germinal centre (GC), proximal follicular B cell marginal zone (FoB-MZ), and marginal zone (MZ). Five GCs were assessed, each with triplicate line scans. *p*=0.017 for GC cells compared to T cells, and *p*=0.044 for FoB-MZ cells compared to T cells, Student’s T-test.

### Dissipation of CIZ1-Xi assemblies during a window in lymphocyte differentiation

As reported for *Xist*, Xi-associated CIZ1 protein assemblies are not detected in naive B and T cells but become re-enriched in the Xi territory upon stimulation with Lipopolysaccharide (LPS) or αCD3, which trigger B and T cell activation respectively (Fig.2C,D Supplemental Fig.1D) (Ridings-Figueroa et al., 2017). Here, immunodetection of CIZ1 allowed the frequency of CIZ1-Xi assemblies to be quantified after 24 hours, when a major transition in B cell gene expression occurs (Shi et al., 2015), and again at 48 hours (Supplemental Fig.1E), and their classification into three categories: nuclei with no CIZ1-Xi assembly, nuclei with discrete apparently fully formed assemblies, and nuclei with intermediate ‘dispersed’ assemblies (Fig.2E). The proportion of cells with fully formed CIZ1-Xi assemblies was low in the naive populations, nearly doubled for B cells and more than tripled for T cells within the first 24-hour window.

Upstream in the haematopoietic lineage cascade, bone marrow-derived long-term haematopoietic stem cells (LT-HSCs) (Kent et al., 2009), short-term HSCs (ST-HSCs), and multipotent-progenitors with lymphoid lineage potential (MPP-Ly) sorted by flow cytometry (Fig.2F,G), presented discrete CIZ1-Xi assemblies in 77%, 82%, and 85% of their populations, respectively (Fig.2E,H). Thus, CIZ1 is enriched around Xi chromatin in stem and lineage restricted cells, dissipates during the prolonged quiescence of mature naive B and T lymphocytes, and then condenses around the Xi again upon activation by antigen. This description of CIZ1 behaviour aligns with *Xist* behaviour in the haematopoietic lineage (Grigoryan et al., 2021; Savarese et al., 2006), and confirms the existence of a prolonged window in B cell differentiation in which the Xi is uncloaked.

### CIZ1-Xi assemblies in germinal centre B cells *in vivo*

The correlation between CIZ1 and *Xist* behaviours makes CIZ1 an informative protein marker to explore the spatial dynamics of repressive molecular assemblies at the Xi *in vivo*, where B cells activate in a T cell-dependent B cell receptor-driven process. In whole-spleen cryosections from a young WT female the pan-B cell marker B220 was used to identify B cells, Enhancer of Zeste Homolog 2 (EZH2) to mark germinal centre sites of B cell activation and expansion (Beguelin et al., 2013), and Siglec-1 (CD169/MOMA1) to mark metallophillic macrophages at the boundary between splenic red and white pulp (Fig.2I). Together these demarcate five functional zones across the tissue; the T cell zone (B220^−^ EZH2^−^), follicular B cell zone (FoB, B220^+^ EZH2^−^), germinal centre (GC, B220^+^ EZH2^+^), proximal follicular B cell marginal zone (FoB-MZ, B220^+^, EZH2^−^ Siglec-1^lo^) and the marginal zone (MZ, B220^lo^ EZH2^−^ Siglec-1^+^). A significant increase in discrete CIZ1-Xi assemblies is evident within the GC, which decreases as B cells become more proximal to the MZ (Fig.2I). Thus, the CIZ1-Xi assembly re-formation quantified in activated B cell populations *in vitro* can be observed *in situ* in germinal centres.

### CIZ1-null lymphocytes respond to LPS activation

We hypothesised that the dissolution of CIZ1-*Xist* assemblies that occurs in naive B and T lymphocytes represents a ‘window of vulnerability’ during their differentiation, in which dosage compensation could be compromised. Furthermore, since CIZ1-null mice cannot form CIZ1-*Xist* assemblies at the Xi, compromised dosage compensation could be exacerbated before, during or after the window, or during the transition each time they face immunological challenge.

We first compared the response to LPS stimulation of WT and CIZ1-null whole spleen-derived cell populations from 8–10 week old female mice and observed no gross differences, with both showing patches of clonal expansion after 24 hours (Fig.3A,B) and increased expression of EZH2 and mutator enzyme AID (*Aicda*) responsible for driving somatic hypermutation and class-switch recombination (Muramatsu et al., 2000) (Supplemental Fig.2A). Analysis of transcription factor gene signatures associated with stages of B cell maturation, including 43 genes associated with naive B cells, and 23 germinal centre genes (Shi et al., 2015), confirmed that WT and CIZ1-null cell populations behaved similarly by dampening the naive signature, and elevating the activated germinal centre signature (Fig.3C, Supplemental Data set 2 and 3). Thus, the initial response to activation appears to be largely intact in the absence of CIZ1.

**Fig. 3.**
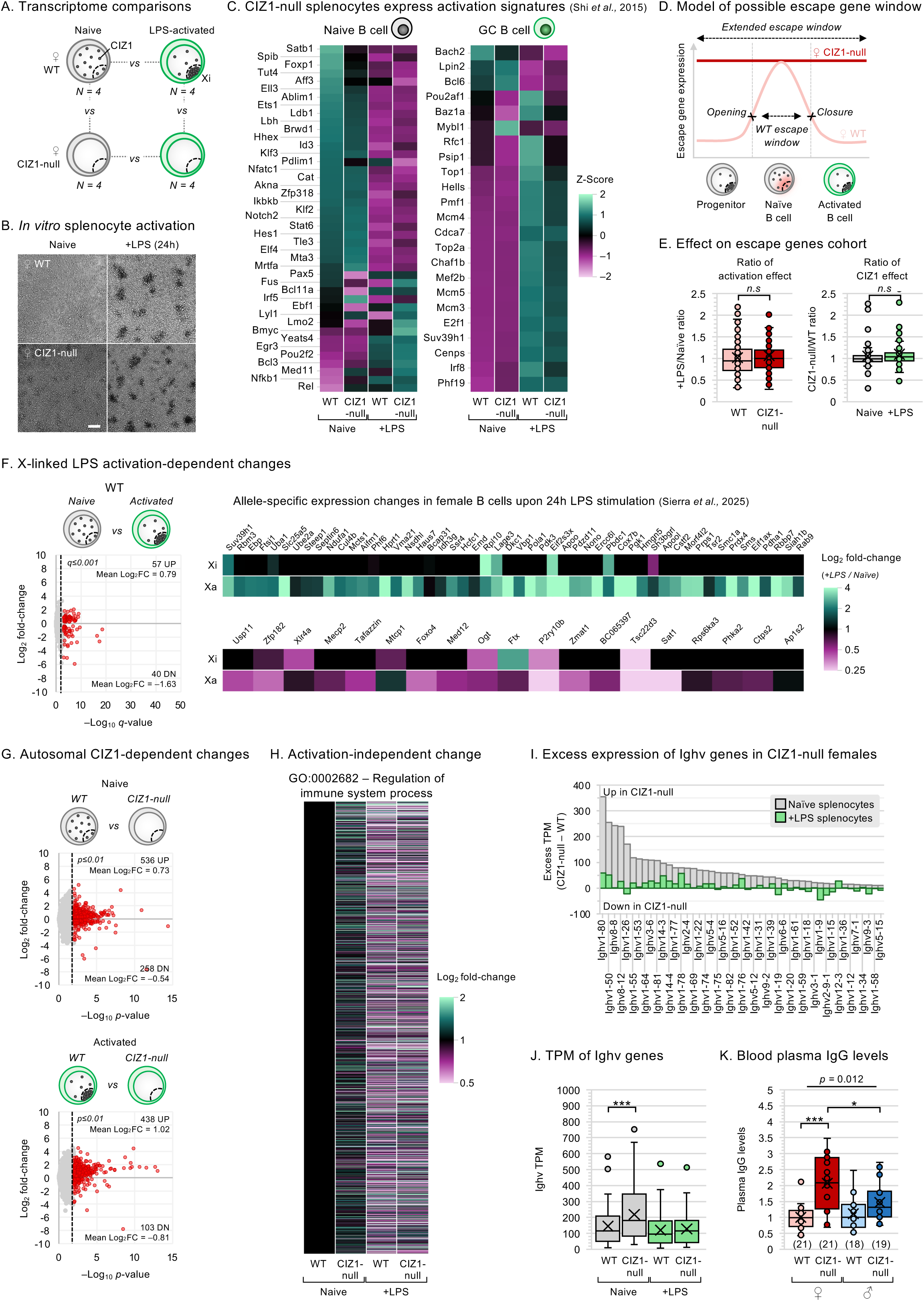
Premature activation-independent changes in female CIZ1-null naive lymphocytes A) Schematic showing transcriptome comparisons between WT and CIZ1-null female naive and 24-hour LPS-activated splenocyte populations, based on four replicates of each genotype and state. B cell activation-dependent changes (with and without CIZ1), and state-specific differences are derived. B) Brightfield images of female WT and CIZ1-null total splenocytes following LPS activation for 24 hours. Scale bar is 100 µm. C) Heatmaps showing Z-scores for transcription factor gene signatures associated with B cell activation and maturation (Shi et al., 2015). GC = germinal centre. TPM average and standard deviation across all samples was calculated per gene to determine Z-score. D) Window of vulnerability model depicting escape gene expression before and after CIZ1-*Xist* assembly dispersal from the Xi in B cell development, with elevated escape gene expression throughout in the absence of CIZ1. E) TPM ratios of the 133 escape gene (Hauth et al., 2026) in female splenocytes shown as activation effect in WT and CIZ1-null (left; LPS-activated vs naive) and CIZ1 effect in naive and LPS-activated populations (right; CIZ1-null vs WT). F) (Left) Volcano plot showing activation-dependent changes in X-linked genes (*q*≤0.001) in WT female splenocytes from 8–10 week old mice. (Right) Heatmap depicting allele-specific log_2_ fold-changes of X-linked activation-dependent DEGs in Sierra et al. (2025) dataset for 52/57 found UP DEGs and 19/40 found DN DEGs across three internal replicates. G) Volcano plots showing CIZ1-dependent changes of autosomal genes (*p*≤0.01) in female splenocytes from 8–10 week old mice, in naive (top) and LPS-activated (bottom) states. H) Heatmap showing TPM log_2_ fold-change for the 1689/1759 genes within the gene signature ‘Regulation of immune system process’ (GO:0002682). Genes that had TPM = 0 across all four replicates (70 genes) were removed. Average TPM across four replicates were normalised to WT naive female splenocytes. I) Excess TPM in CIZ1-null over WT for female naive splenocytes (grey) and +LPS splenocytes (green), showing 47/113 Ighv genes with a CIZ1-null excess TPM ≥ 10 in naive splenocytes. J) TPM levels of the 47 Ighv genes in female WT and CIZ1-null naive (grey) and +LPS (green) splenocytes. K) IgG protein levels in blood plasma quantified by ELISA, normalised to mean of female WT. Number of mice are shown in parentheses. Analysis by Two-Way ANOVA.

However, globally the gene expression changes observed in CIZ1-null cells upon LPS stimulation differ from WT, because less than half the number of genes meet the significance threshold (FDR *q*≤0.001, Supplemental Fig.2B). Most of the CIZ1-null transition DEGs are shared with WT (85/77% of autosomal UP/DN DEGs), but 2326 are ‘missing’ (Supplemental Data set 3). In fact, this cohort undergoes similar but smaller changes, with greater variance amongst replicates (Supplemental Fig.2C,D), suggesting that CIZ1 contributes to the stability of transcriptional responses in female lymphocytes.

### No evidence for window of escape gene expression

If dissolution of *Xist* and CIZ1 from the Xi in WT naive female lymphocytes creates a window of vulnerability for Xi escape, it follows that their re-assembly at the Xi upon activation would later dampen expression, so defining window closure. In CIZ1-null cells, in which the Xi is not populated by CIZ1 assemblies rendering *Xist* unable to accumulate in the Xi territory (Ridings-Figueroa et al., 2017), the escape window may never close (Fig.3D).

We first tested window closure in WT cells by looking at the expression of a set of 133 murine genes that are reported to escape Xi repression in one or more cell type (Hauth et al., 2026). Across the cohort, we observed no overall reduction in expression 24 hours after stimulation of WT lymphocytes with LPS, even though CIZ1-*Xist* assemblies have begun to re-assemble in the Xi territory in this time (Fig.3E and Fig.2E). Similarly, no shift in this cohort was observed in CIZ1-null cells, or between WT and CIZ1-null cells (Fig.3E). Moreover, although we do observe significant changes in X-linked gene expression upon lymphocyte activation in WT (57 UP and 40 DN), an analysis of their expression in published allele-specific transcript data (Sierra et al., 2025), suggests that these are predominantly changing expression from the Xa (Fig.3F). Overall, this data does not support the idea of an Xi escape window that closes in WT female lymphocytes, or an extended window in CIZ1-null female lymphocytes.

### Changes in CIZ1-null lymphocytes prior to activation

Notably, the difference between WT and CIZ1-null lymphocytes in the naive state (794 autosomal DEGs at *p*≤0.01), is greater than in the LPS-stimulated state (541 autosomal DEGs, Fig.3G), with a similar picture on the X chromosome (Supplemental Fig.2E, Supplemental Data sets 4 and 5).

GSEA of CIZ1-dependent UP DEGs showed immune-related and cell activation signatures after LPS-stimulation, but notably these are also evident in the naive state prior to stimulation with LPS (Supplemental Fig.2F, Supplemental Data sets 4 and 5). Moreover, the most significant mutual gene signature (Regulation of immune system process; GO:0002682) is globally elevated in naive CIZ1-null lymphocytes (Fig.3H), and is accompanied by increased expression of immunoglobulin heavy-chain variable region (Ighv) genes (Fig.3I,J). Consistent with this, IgG levels in blood plasma from non-immunized animals were significantly elevated in CIZ1-null females, and this was not seen in males (Fig.3K).

Together, these data illustrate a premature shift toward immune cell activation in females in the absence of challenge. This is before the point at which CIZ1 and *Xist* would normally re-assemble around Xi chromatin and so argues against reformation of *Xist*-dependent assemblies as a primary point of failure underlying CIZ1 pathology.

A similar message emerges in independent data derived from *Xist-*deleted female naive and stimulated B cells (Sierra et al., 2025), which revealed a greater number of differentially expressed genes in the naive state compared to the stimulated state (Sierra et al., 2025). We conclude that, although CIZ1-null female lymphocytes are capable of responding, they have diverged prior to stimulation and acquired a ‘pre-activated’ state.

### Sexually divergent response to absence of CIZ1 in naive B cells

Analysis of lymphocytes before and after stimulation shifted focus away from a potentially faulty transition and inaccurate endpoint, and on to the resting state of naive cells. To probe this more specifically, naive B cells from WT and CIZ1-null animals were enriched from total splenocyte populations using MACS negative-selection, purifying working populations from an average of 47% B220^+^ cells to 90% (Supplemental Fig.3A,B). This allowed a focused analysis of the B cell transcriptome, and was combined with a comparison between the sexes.

In CIZ1-null animals, dampening of the naive B cell transcription factor signature was confirmed and evident in both sexes, while a striking elevation of germinal centre transcription factor signature occurred in females only (Fig.4A, Supplemental Data set 6). Moreover, female naive B cells showed significant elevation of *Aicda* transcripts whereas males did not (Supplemental Fig.3C). This sexually dimorphic profile shows that, while both male and female naive B cells are sensitive to the absence of CIZ1, their response differs. Moreover, the origin of the divergence lies before or during the window in which the Xi is normally uncloaked.

**Fig. 4.**
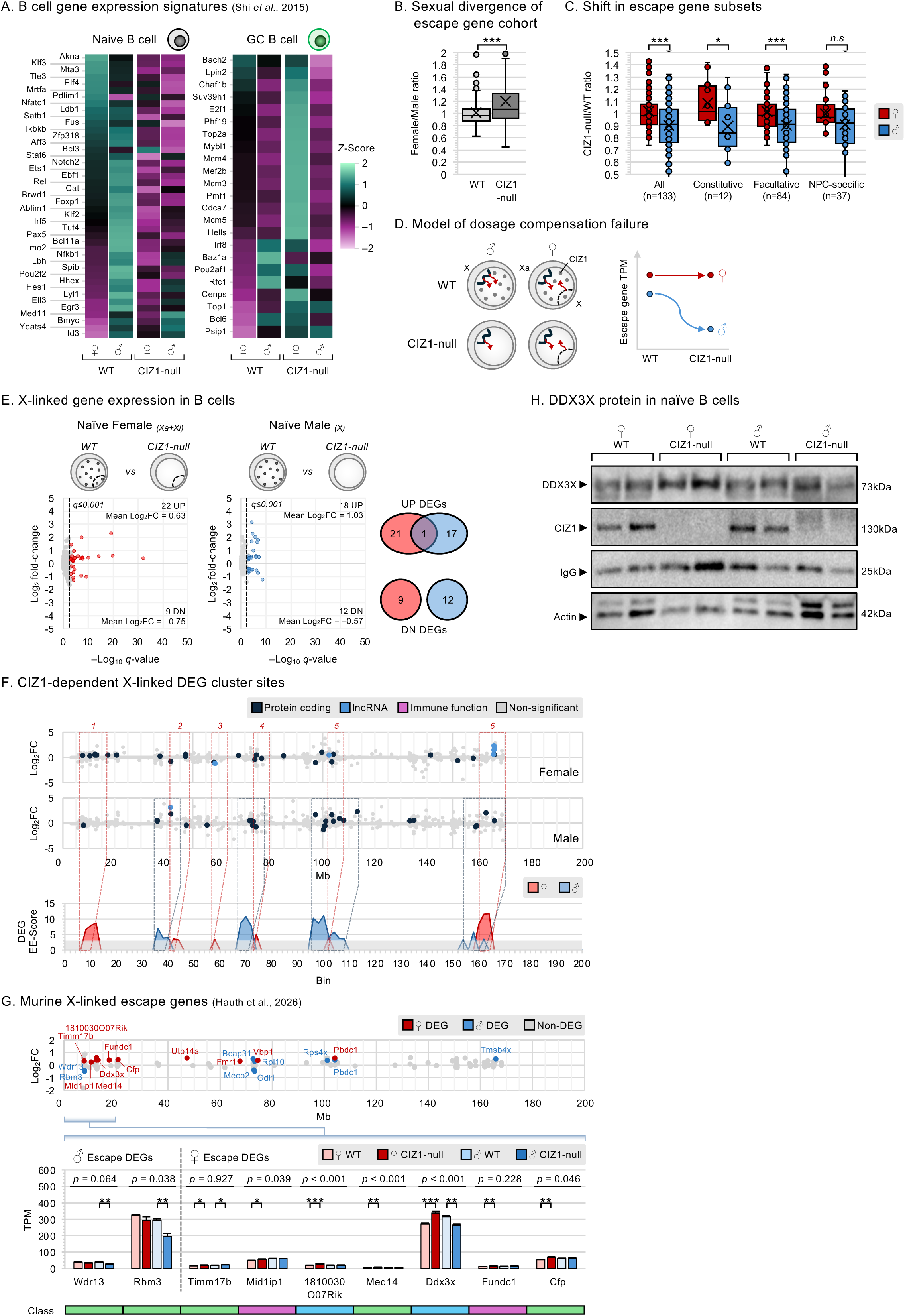
Elevated escape gene expression in female naive B cells A) Heatmaps showing Z-scores in purified B cells for naive and germinal centre transcription factor gene signatures (Shi et al., 2015). B) TPM ratios of the 133 escape genes (Hauth et al., 2026) in WT and CIZ1-null naive B cells shown as female vs male. Analysis by Student’s T-test. C) TPM ratios of the 133 escape genes (Hauth et al., 2026) in naive B cells shown as CIZ1-null vs WT for females (red) and males (blue) split by escape gene subcategory: constitutive, facultative and NPC-specific. Analysis by Student’s T-test, n = number of genes in each subcategory. D) Model of dosage compensation failure from the X chromosome in males, suppressing escape gene expression in the absence of CIZ1. E) Volcano plots showing differentially expressed X-linked genes (*q*≤0.001), comparing 8–10 week old CIZ1-null to WT naive B cells for females (left, red) and males (right, blue). Number of mutual DEGs between the sexes in both UP and DN DEG populations are shown in the Venn diagrams. F) Chromosomal location of X-linked differentially expressed genes in naive B cells, for females and males. Below, gene clusters are defined by an inclusion threshold of EE-Score≥3, showing locations in female (red) and male (blue), and cluster site designations 1-6 above. G) (Top) Chromosomal location of 133 X-linked escape genes (Hauth et al., 2026), highlighting those that are CIZ1-dependent in females (red) and males (blue). (Bottom) Average TPM for CIZ1-dependent escape genes in cluster 1, for female and male DEGs. Asterisks indicate significance between WT and CIZ1-null TPM replicates for females and males by Two-Way ANOVA. Below, reported escape gene classification (Hauth et al., 2026), showing constitutive (blue), facultative (green), NPC-specific (pink). H) Western blot showing DEAD box helicase DDX3X protein, CIZ1, IgG and β-actin loading control in whole cell lysates from naive B cells.

### X-linked escape gene expression in naive B cells

Lack of evidence for escape gene window closure in females, coupled with sexually dimorphic response to lack of CIZ1 in naive B cells shifted focus towards comparisons of X-linked escape genes in males and females. Averaged across the cohort of 133 escape genes (Hauth et al., 2026) expression appears to be dosage compensated, with an expression ratio average close to 1. However, in the absence of CIZ1 the ratio shifts towards more expression in females than males, indicating compromised dosage compensation (Fig.4B). Unexpectedly, this compensation failure is driven by reduced expression in males rather than increased expression in females. While females maintain an average gene expression ratio close to 1 across the cohort, males repress escape genes by an average of 10% across all escape gene subcategories (Fig.4C). Our dataset does not allow female expression to be identified as Xa or Xi-derived, so it remains an open question whether absence of CIZ1 results in Xi effects that are masked by Xa effects (or vice versa). However, it is clear that males experience changes in escape gene expression from their X chromosome when CIZ1 is absent, causing them to diverge from females. Furthermore, this argues for a non-canonical mechanism of dosage compensation that modulates expression in males, and which involves CIZ1 (Fig.4D).

### CIZ1-dependent escape genes

We next asked about specific escape genes and the extent to which they intersect with CIZ1-dependent DEGs. On the X chromosome absence of CIZ1 leads to 31 DEGs in female naive B cells, with twice as many UP than DN, and these define six CIZ1-dependent cluster sites with EE-Score≥3 (Fig.4E,F, Supplemental Data set 6). However, a similar number of genes are affected in males (Supplemental Data set 7), and these define four of the same sites, despite only one of the genes (constitutive escape gene *Pbdc1* at 104,123,362 Mb) undergoing a similar directional change in both sexes. Since any X-linked genes sensitive to CIZ1 absence in males must report on non-Xi mechanisms, these data further indicate influence that is independent of *Xist.* Moreover, as *Pbdc1* is de-repressed in both sexes and to similar levels (Supplemental Fig.3C,D), these data suggest it is better characterised as a CIZ1-dependent gene rather than an *Xist*-dependent gene.

Of the female CIZ1-dependent UP DEGs, 50% (11/22) are escape genes while none are among the female DN DEGs. For males, 28% (5/18) of the UP, and 33% (4/12) of the DN DEGs, are designated Xi escape genes in females (Supplemental Fig.3D and Fig.4G, upper and Supplemental Data sets 6 and 7).

Of the 10 genes that are both escape genes and female-specific CIZ1-dependent UP DEGs (excluding *Pbdc1*), six (*Timm17b*, *Mid1ip1*, *1810030O07Rik*, *Med14*, *Ddx3x*, and *Fundc1*) define female-specific cluster 1 (at 8–18 Mb) and an additional escape gene (*Cfp*) lies just beyond the boundary of our cluster analysis (Fig.4G, lower). This coincides with an independent study that identified disproportionate escape gene expression in WT naive and stimulated B cells in the proximal ∼20Mb region of the Xi (Sierra et al., 2025).

In our study, the DEAD-box RNA helicase gene *Ddx3x* is by far the most highly expressed and also most affected by the absence of CIZ1 in both sexes, with TPMs that are 24% higher in female CIZ1-null naive B cells, but 16% lower in males (Fig.4G, lower). Sex-specific *Ddx3x* transcript level differences are mirrored at the protein level (Fig.4H).

This identifies an escape gene domain that is under CIZ1 control in female B cells (Xi, Xa or both), and highlights DDX3X as a gene product that experiences heightened female-male divergence in the absence of CIZ1. DDX3X has already emerged as an essential factor in B cell development, supporting both proliferative and epigenetic changes necessary for the rearrangement of immunoglobulin genes (Liu et al., 2025). Therefore, DDX3X production in CIZ1-null female naive B cells is a candidate contributor to the B cell lymphoma phenotype.

### Divergent response of autosomal genes in males and females

The effects of CIZ1 absence are not likely to be limited to the X chromosome, as it also populates smaller assemblies throughout the nucleus in both sexes (Fig.5A). Focussing on autosomal genes, absence of CIZ1 leads to 366 UP and 431 DN DEGs in female naive B cells, and 471 UP and 379 DN DEGs in males (*q*≤0.001) (Fig.5B, Supplemental Data sets 6 and 7). However, less than 10% are shared, and of the female UP and DN DEGs, 340/366 (93%) and 281/431 (65%) remain unchanged in males, with a similar reciprocal picture (Fig.5C).

**Fig. 5.**
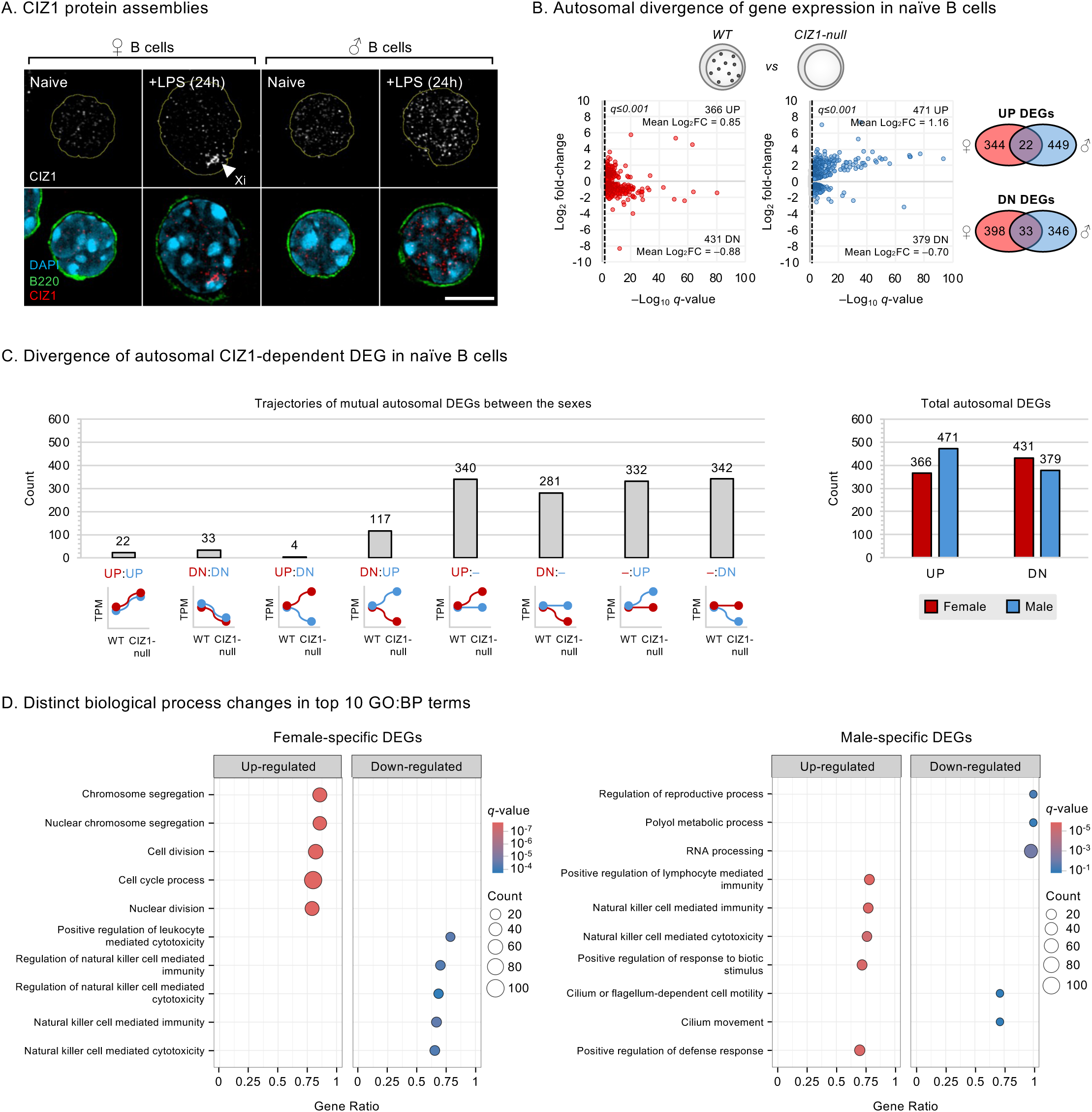
Sex-specific response of autosomal genes to CIZ1 absence A) CIZ1 protein assemblies in female and male naive and LPS-activated B220^+^ B cell nuclei. Images were acquired as Z-stacks by super-resolution microscopy, and CIZ1 (red) and DAPI (blue) channels flattened. B220 (green) is shown as a single central slice in the merged images. Above, CIZ1 in grey scale showing Xi and non-Xi assemblies. Scale bar is 5 µm. B) Volcano plots showing autosomal CIZ1-dependent changes (*q*≤0.001) in 8–10 week old naive B cells for female (left, red) and male (right, blue). Venn diagrams show mutual UP and DN DEGs between the sexes. C) Left, comparison of trajectories of mutual autosomal DEGs between females (red) and males (blue). Right, bar plot showing distribution of total UP and DN regulated autosomal CIZ1-dependent DEGs. D) The top five UP and DN gene expression pathways (GSEA M5 biological processes) affected by the absence of CIZ1 in naive B cells (autosomal and X-linked DEGs combined) for female-specific DEGs (left) and male-specific DEGs (right).

Crucially, some genes show divergent changes in expression, where their response to the absence of CIZ1 is significantly opposite between sexes – 117 DEGs significantly decrease in females but increase in males (four do the reverse) (Fig.5C). This emphasises the dampening effect that CIZ1 has on the polarisation of gene expression. Overall, the autosomal profile conflicts with what might be expected from extrapolation from the X chromosome in that absence of CIZ1 is not associated with de-repression specifically, but with change.

### De-repression of female-specific autosomal genes promotes cell proliferation

Since the majority of CIZ1-dependent genes diverge in males and females the absence of CIZ1 would be expected to promote sex-specific cell identities, and indeed GSEA of M5 Gene Ontology Biological Process revealed increased expression of pro-proliferation gene sets in females that was not seen in males (Fig.5D, Supplemental Fig.3E), as well as dampening of natural killer cell-associated gene sets. Conversely, the gene sets enriched in males promote immune cell function (Fig.5D). These divergent gene signatures indicate that CIZ1-null males are as likely as females to experience aberrant gene expression and altered identity. However, de-repression of pro-proliferation genes likely underpins the sex-specific B cell lymphoma phenotype that emerges exclusively in CIZ1-null females.

### Gene clusters and gene families

Cluster analysis of autosomal B cell DEGs identified 64 EE-Score-defined sites in females and 56 in males across all chromosomes, of which 34 occur in overlapping genomic regions (Fig.6A, Supplemental Fig.4A–C, Supplemental Data sets 6–8). In some cases, the DEGs that define common cluster sites are different, and in others they are common to both males and females but do not respond in the same way. At least three gene families that define immune cell phenotype experience polarised expression in the absence of CIZ1 (Fig.6B,C). These include the cluster site at 130Mb on chromosome 6 which encodes 19 Killer cell Lectin-like Receptor (*KLR*) genes, 18 of which increase in males but decrease in females to the extent that the domain is defined as a CIZ1-dependent site in both sexes based on EE-score. Divergent expression was tested at the protein level in whole cell lysates from naive B cells using two independent antibodies to KLR family members, which both detected a CIZ1-dependent protein at 70kDa (Supplemental Fig.4D). Similarly, the *Ighg/Ighv* genes located at 110–122 Mb on chromosome 12 also show opposing responses between the sexes, that also contrasts with the KLR gene cluster by increasing in females and decreasing in males. Other locations meet EE-score criteria in only one sex but display polarising trends. For example, the T cell receptor beta (*TCRβ*) locus at 40Mb on chromosome 6 harbours variable and joining (*Trbj/Trbv*) genes, of which some decrease significantly in females enough to define a cluster, while in males the threshold is not met (Fig.6B,C).

**Fig. 6.**
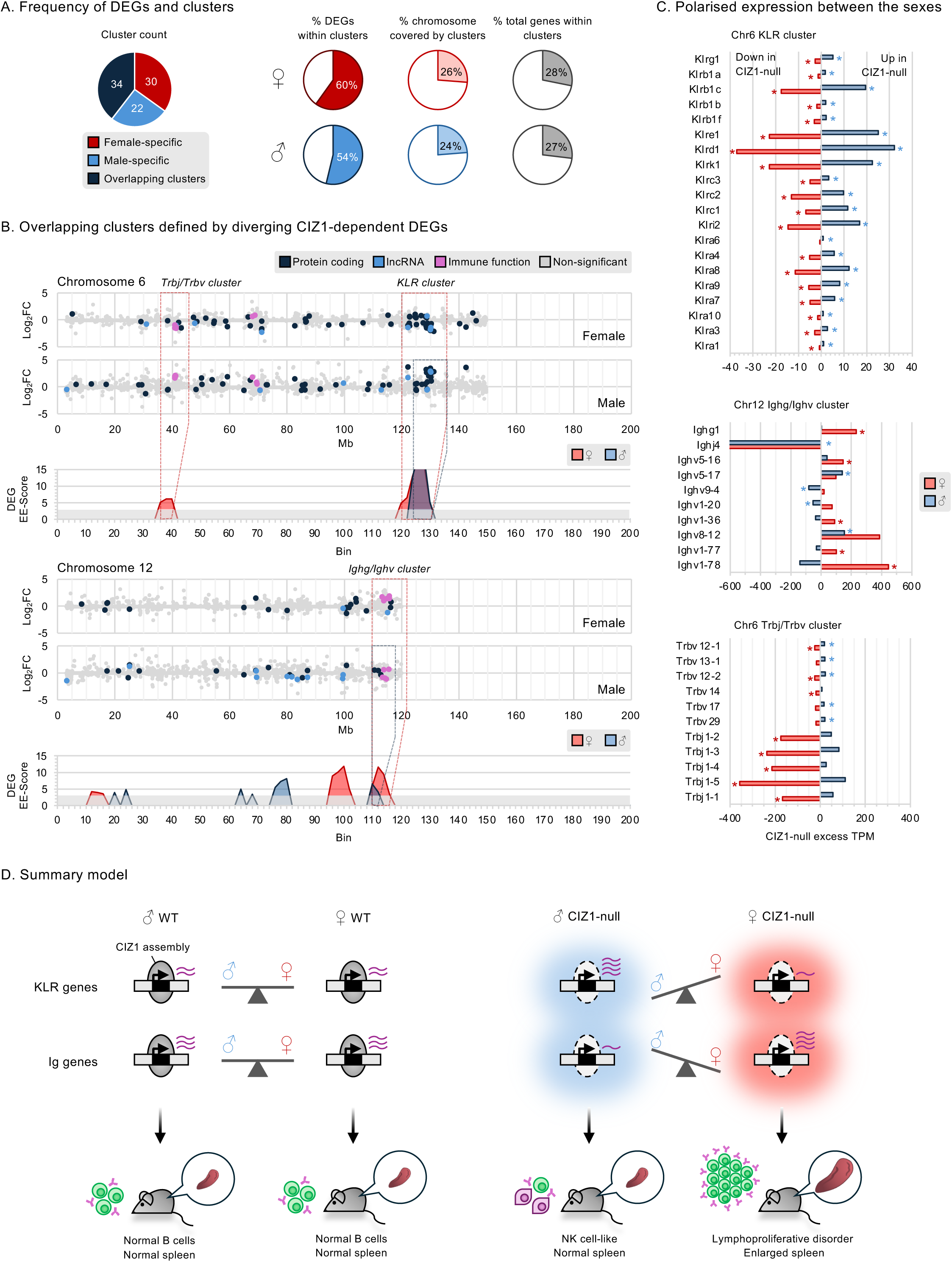
Divergent immune gene clusters and polarisation of gene expression between sexes A) Pie chart shows average distribution of CIZ1-dependent DEG-defined clusters in female (red) and male (blue) naive B cells, and overlapping sites between the sexes (dark blue). Right, pie charts showing average proportions of DEGs within cluster sites (left), chromosome coverage of cluster sites (EE-Score≥3, centre), and total genes found within clusters (EE-Score≥3, right), for female and male WT vs CIZ1-null comparisons. B) Location of DEGs in female and male naive B cells for chromosomes 6 and 12. Below, EE-Score≥3 derived gene cluster sites for female (red) and male (blue), with overlapping clusters of gene families that exhibit polarised expression between sexes outlined. C) Excess TPMs in CIZ1-null over WT for genes in overlapping cluster families, showing females (red) and males (blue). Asterisks denote whether individual genes are CIZ1-dependent DEGs (*q*≤0.001). D) Model of CIZ1 modulating the divergent response of autosomal gene clusters in female and male naive B cells to drive phenotypic divergence.

These data show that divergent transcriptional responses to absence of CIZ1 can occur at common locations, sometimes affecting the same genes, and that those identified by the statistical thresholding applied here are unlikely to be the whole story. Overall, the picture that emerges suggests that CIZ1 assemblies may form a spatially confined environment that modulates and fine-tunes the response of selected chromosomal regions to sex-specific influences.

## DISCUSSION

Integration of compromised X-linked escape gene dosage compensation with the polarised response of some autosomal gene clusters, leads us to propose a unifying model in which the dual effects interact to give rise to sex-biased immune system dysfunction in CIZ1-null mice. First, absence of CIZ1-RNA protein assemblies affects underlying chromatin across large autosomal domains in both sexes. Our previous analysis of Xi chromatin in female fibroblasts showed that transient disruption of CIZ1 assemblies leads to rapid loss of H2AK119Ub in a deubiquitylase (DUB)-dependent manner, suggesting that CIZ1 assemblies normally restrict access of polycomb repressive complex 1 (PRC1)-DUBs to underlying Xi chromatin (Turvey et al., 2025). Extrapolation to other enzymatic or regulatory functions would suggest that de-protected loci become vulnerable to the prevailing cohort of diffusible molecules in the cell.

Second, failure to dosage compensate X-linked escape gene expression shifts the prevailing cohort of diffusible molecules towards an exaggerated state. Unexpectedly, this involves changes in male X-lined gene expression, rather than female. Together, these dual functions suggest that CIZ1 acts to dampen polarisation between sexes, by protecting common autosomal sites in males and females from sexually divergent influences (Fig.6D).

A candidate gene product that could contribute consequential sexually divergent effects is DDX3X because, within the CIZ1-dependent escape gene cluster at 8–18 Mb, it is by far the most highly expressed and also most affected by absence of CIZ1 (in both sexes).

In fact, DDX3X is already implicated in a diverse range of sex-biased diseases encompassing intellectual disability disorders and cancer (Gadek et al., 2023). Like its Y-linked paralog DDX3Y, DDX3X encodes a DEAD-box RNA helicase that regulates ribonucleoprotein complex assemblies, which are integral to the RNA life cycle and normal development. Both form phase separated molecular condensates, and while they have similar structure, DDX3Y forms more robust condensates than DDX3X suggesting that they may differentially impact cellular function (Shen et al., 2022). Furthermore, DDX3X can supress DDX3Y, and at the protein level they interact with each other implying that the balance between them is a critical factor (Xu and Wei, 2025). DDX3X is known to be required for the normal expansion and function of germinal centre B cells, and *Ddx3x* deletion alters lymphoid development in female mice specifically (Lacroix et al., 2022). Moreover, loss of function mutations in DDX3X are prevalent in human male germinal centre-derived B-cell malignancies, but not females (Lacroix et al., 2023). This complex picture is further nuanced because loss of *DDX3X* function initially helps lymphomagenesis by buffering the proteotoxic stress driven by c-MYC activation, while progression requires DDX3 activity and is achieved in male human primary GC B cells by upregulation of DDX3Y (Gong et al., 2021).

Thus, over-expression of DDX3X in female CIZ1-null cells is expected to be consequential for cell proliferation and cellular identity.

Immune cell composition is typically uneven between the sexes in both humans and mice and can, in extremes, manifest as disease with uneven incidence. Females have greater CD19^+^ B cell numbers and IgM concentration underpinning a bias toward humoral immunity, while males tend toward greater natural killer (NK) cell numbers in the spleen, and their B cells have a tendency toward NK-like gene expression (Cheng et al., 2023; Radovanovic et al., 2026). Thus, the phenotypic consequences for mice lacking CIZ1 appears to be to shift further in the direction of their normal sex-biased tendency. In female CIZ1-null mice we observe more B cells that significantly up-regulate immunoglobulin variable region (IgV) encoding genes, accompanied by a shift towards a germinal centre gene expression profile including *Bcl6* (from a CIZ1-dependent cluster site at 14–28 Mb on Chr 16), and down-regulation of the *KLR* gene family (120–136 Mb cluster on Chr 6), and *Trbj/Trbv* gene family (36–46 Mb cluster on Chr 6). The *KLR* gene family is primarily expressed on NK and T cells acting to recognise MHC class I proteins to regulate immune responses (Raulet, 2003; Yokoyama and Plougastel, 2003), while *Trbj/Trbv* genes are responsible for generating the repertoire of T cell receptors in T cells (Anderson and da Rocha, 2022; Bosc and Lefranc, 2000). In contrast, male naive B cells shift towards NK-like gene expression, elevating *KLR* genes, and do not exhibit a shift towards germinal centre gene expression. Other changes that dampen their B cell identity include down-regulation of *Irf4* which is critical for maturation from pre-B to B cell (Lu et al., 2003), and *Pax5* (CIZ1-dependent cluster 44–54 Mb on Chr 4) which is considered a master regulator of B cell lineage commitment (Medvedovic et al., 2011). Taken together, the gene expression changes show that absence of CIZ1 enables female and male naive B cells to shift in different directions, and that these directions reflect their normal tendency. This argues for a cell intrinsic fate determination, with sex-bias that is influenced by CIZ1.

### Outstanding questions and limitations

This analysis poses some obvious questions. For example, it is notable that several CIZ1-dependent cluster sites encode immune gene families that are rapidly evolving (KLR, BCR, TCR) or susceptible to germline recombination (BCR/TCR). In fact, in the male germline, CIZ1 is considerably elevated in pachytene spermatocytes during the window of recombination (Greaves et al., 2012), raising questions of whether CIZ1 participates in or protects against these processes. It also remains unclear how CIZ1-protected autosomal domains are selected and why they are so large. By analogy with *Xist*-dependence at Xi, and the growing understanding of how RNA-seeded molecular condensates shape the nuclear architecture, our working hypothesis is that localised high-density RNAs seed the formation of CIZ1-containing condensates which then grow and dissipate in a regulated manner (Byrom et al., 2026; Quinodoz et al., 2021). Regarding DDX3X, at present there is also no direct evidence that it functions in concert with CIZ1. While DDX3X has been reported as a protein interaction partner of CIZ1 (Thacker et al., 2020), a functional interaction at the level of CIZ1-RNA assemblies is inferred. Finally, we cannot be definitive about the origin of the escape gene transcripts in females because our genetic model does not support allelic discrimination. However, whether their origin is Xa or Xi in females, CIZ1-null cells experience alterations in the dosage compensation of X-linked escape genes that is driven by changes in males via non-Xi mechanisms, with all of the accompanying potential to affect cell identity.

## Supporting information

Data set 1

Data set 2

Data set 3

Data set 4

Data set 5

Data set 6

Data set 7

Data set 8

## Acknowledgments

We are grateful to Gina Doody and Ulf Klein for guidance on spleen cell markers, to Damian Perez-Mazliah for guidance on B cell isolation, and to Alastair Droop, Andrew Mason and Marcello Beltrami for guidance on transcriptomic analysis, and the Biological Service Facility at the University of York for daily maintenance and welfare checks on our mouse colonies. We would also like to thank Emma Stewart for additional RNA isolation, Zuzanna Bartczak for additional B cell analysis, and colleagues for teaching specialised techniques including Jenny Baker for cryo-sectioning, Naj Brown for tail-vein bleeds, Karen Hogg, Graeme Park and Karen Hodgkinson for flow cytometry and cell sorting, Grant Calder for high-resolution microscopy and Susanna Rose for ELISA assays.

## Competing interests statement

DC reports receiving institutional research support from Cizzle Biotech. DC and JA are founders of Cizzle Biotech. Other authors disclosed no potential conflicts of interest.

## Funding

This work was supported by the Medical Research Council DiMeN PhD studentship to EG (MP/W006944/1), a GenerationResearch summer placement award (N0017316) to RM, and Medical Research Council grant MR/V029088/1 to DC.

## Author contributions

Conceptualisation – EG, DC, JA

Methodology – EG, SJ, FP, JA, DC

Investigation – EG, RM, JP, DC, JA

Software –

Formal analysis – EG, FP

Resources – EG, JP, JH, JA

Writing, review, editing – EG, JH, DK, DC, JA

Funding acquisition – DC

## MATERIALS AND METHODS

### Data availability statement

Further information and requests for resources and reagents should be directed to the lead contacts Elena Guglielmi and Dawn Coverley.

### Key resource table

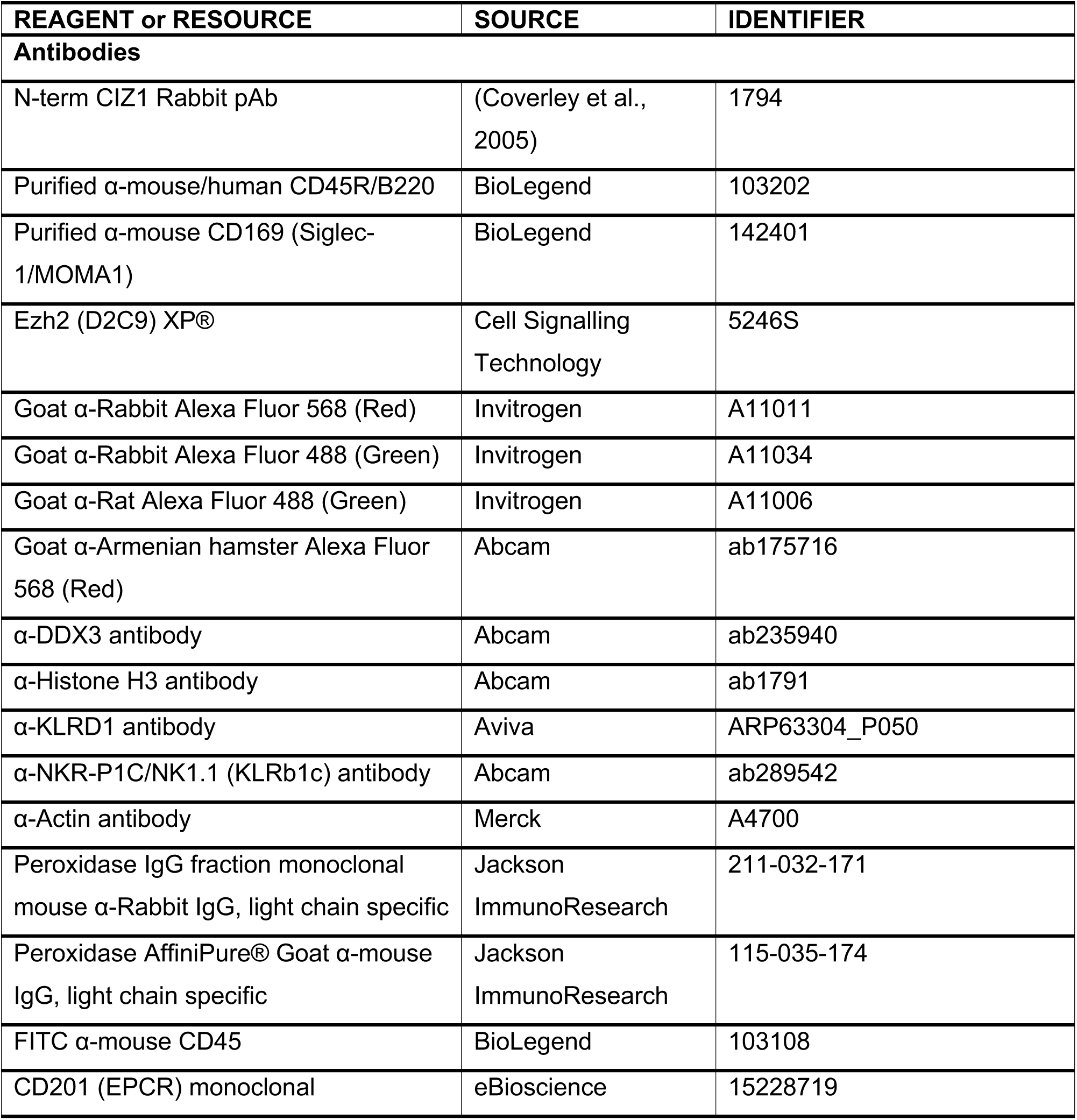
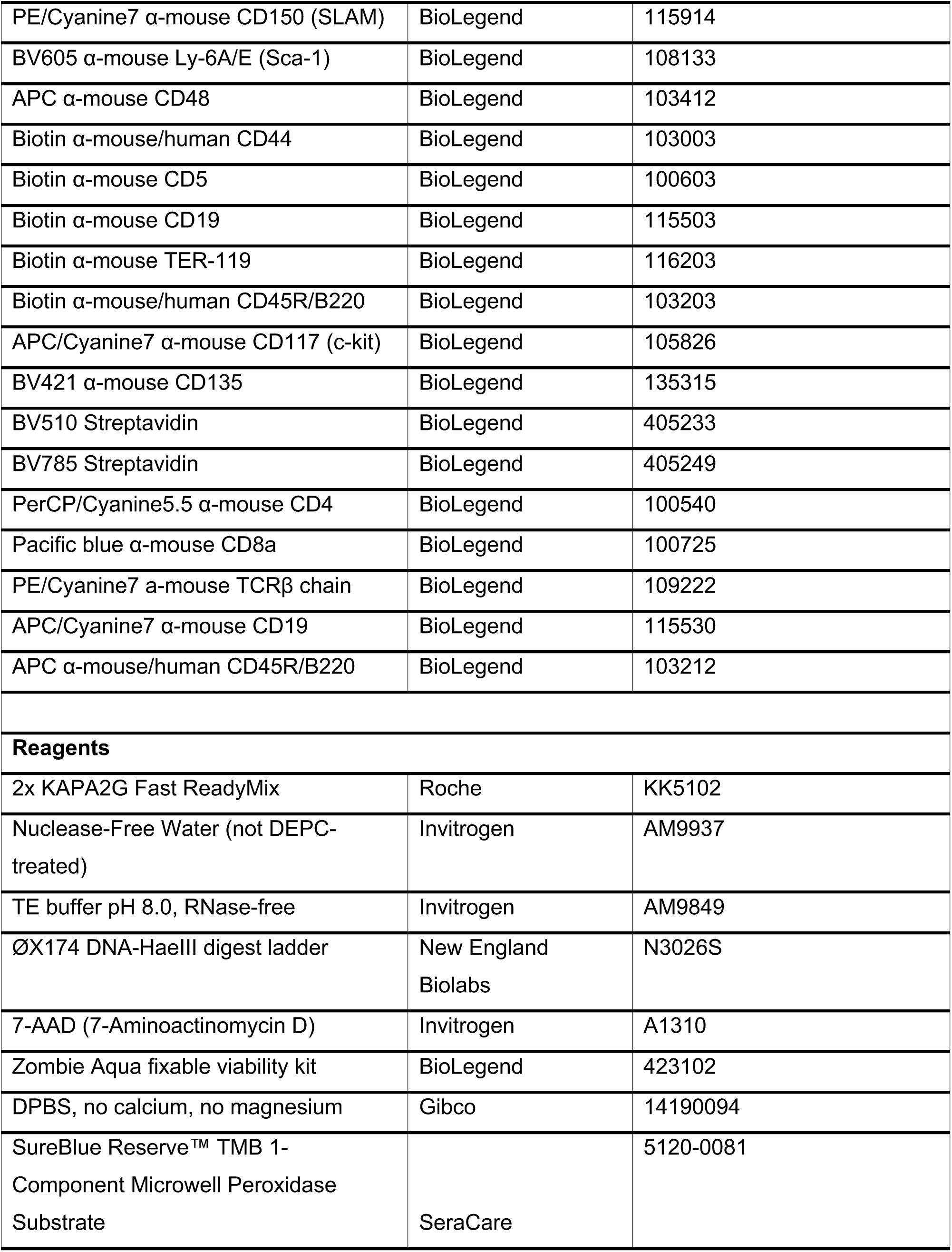
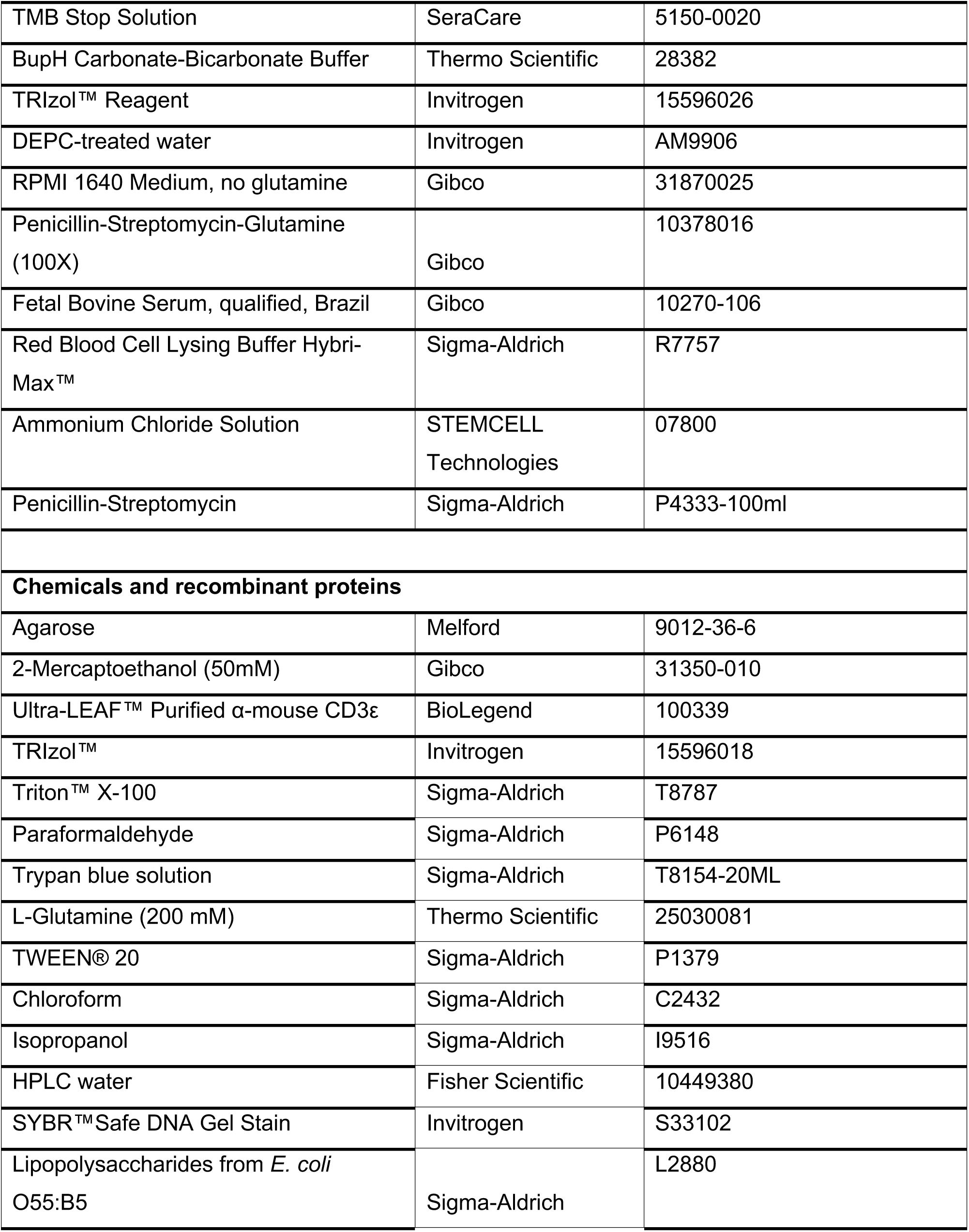
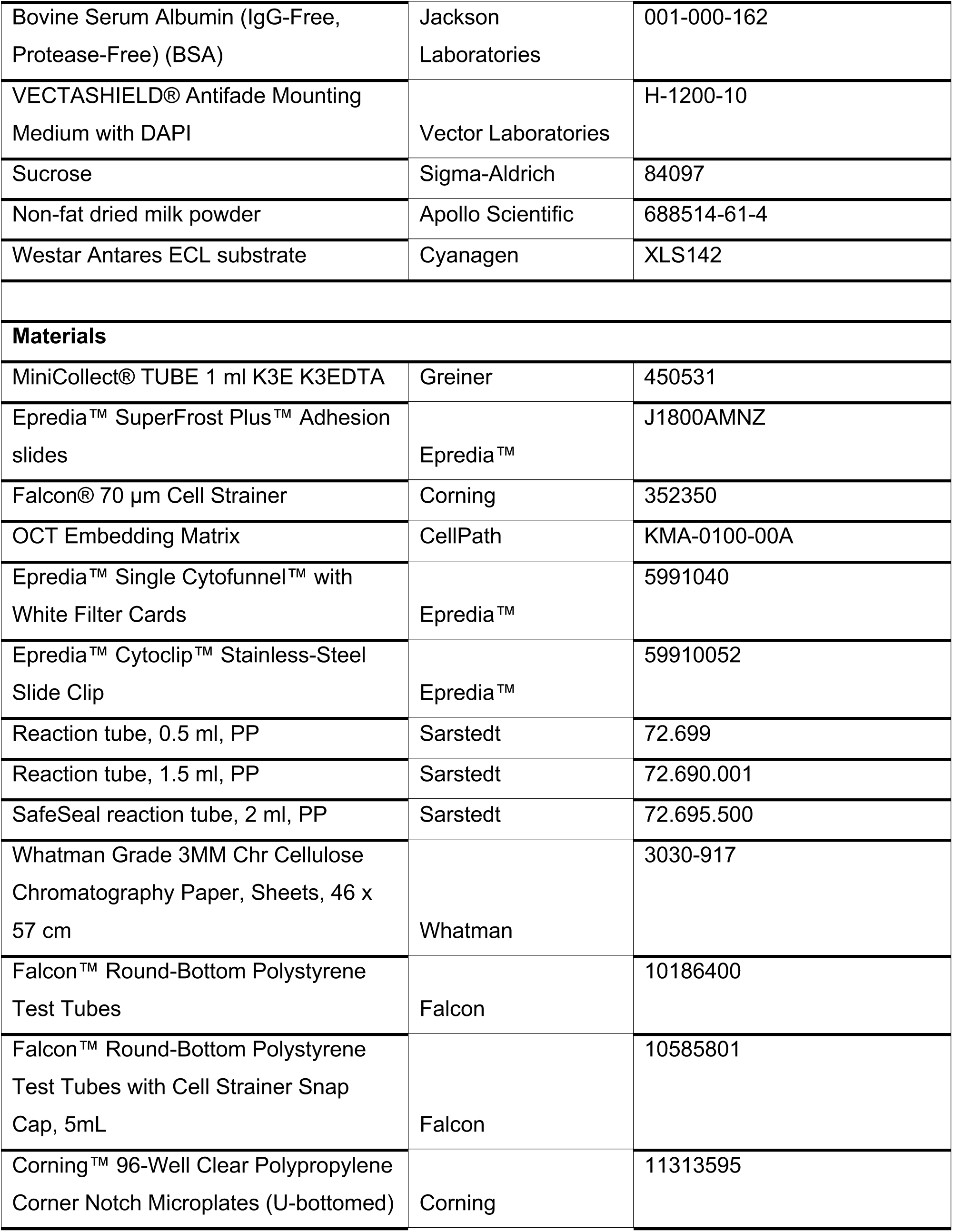
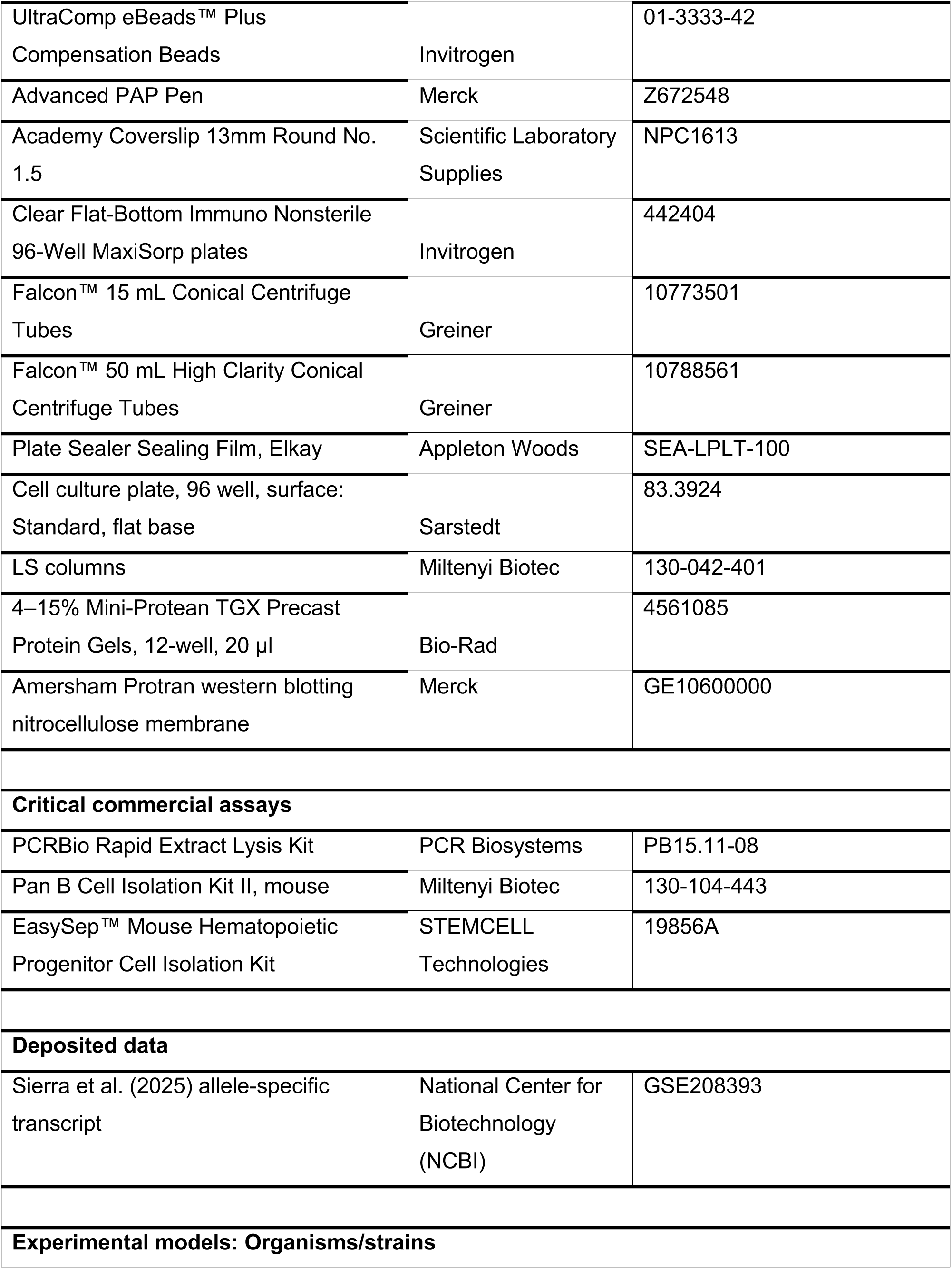
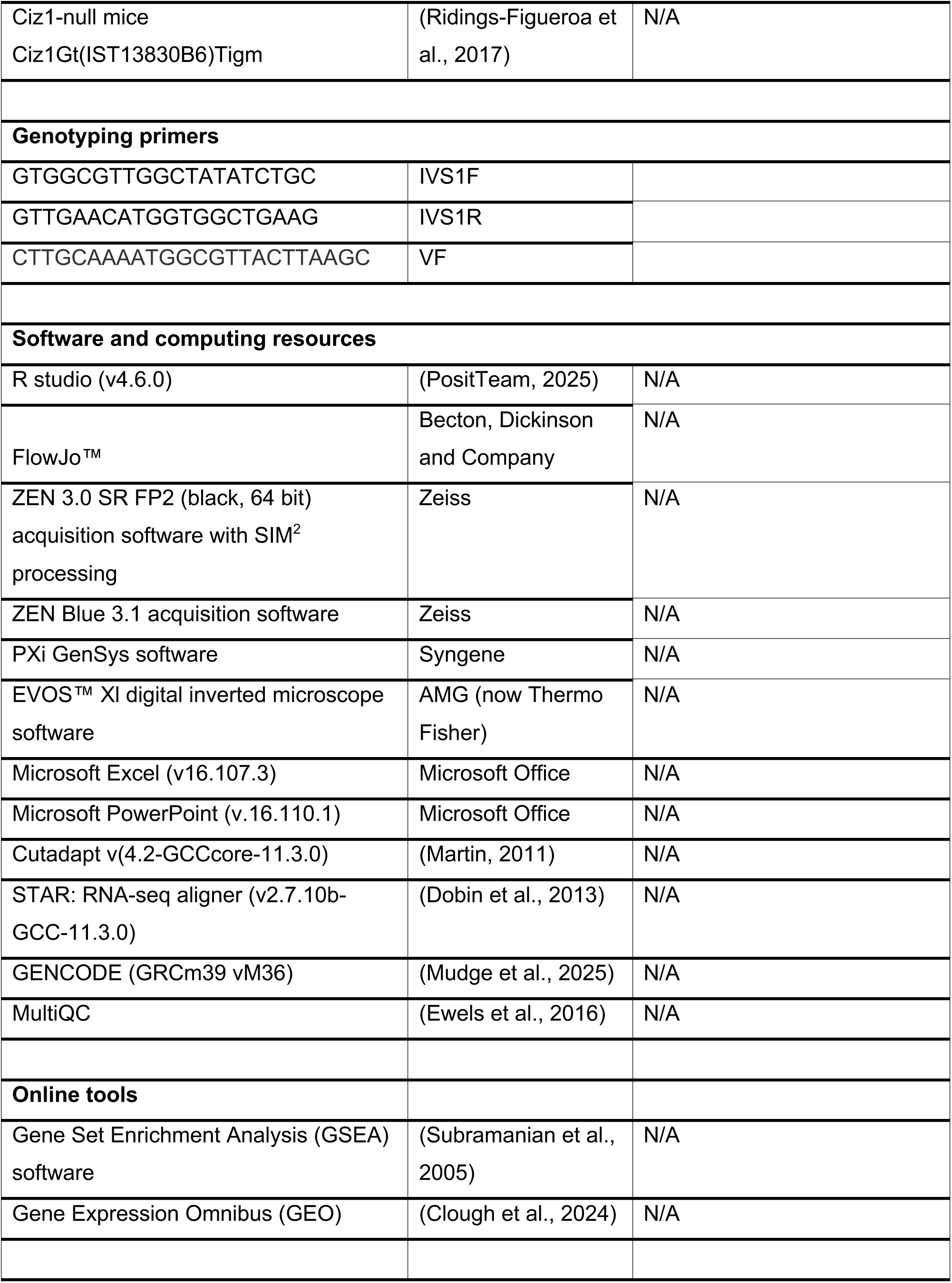
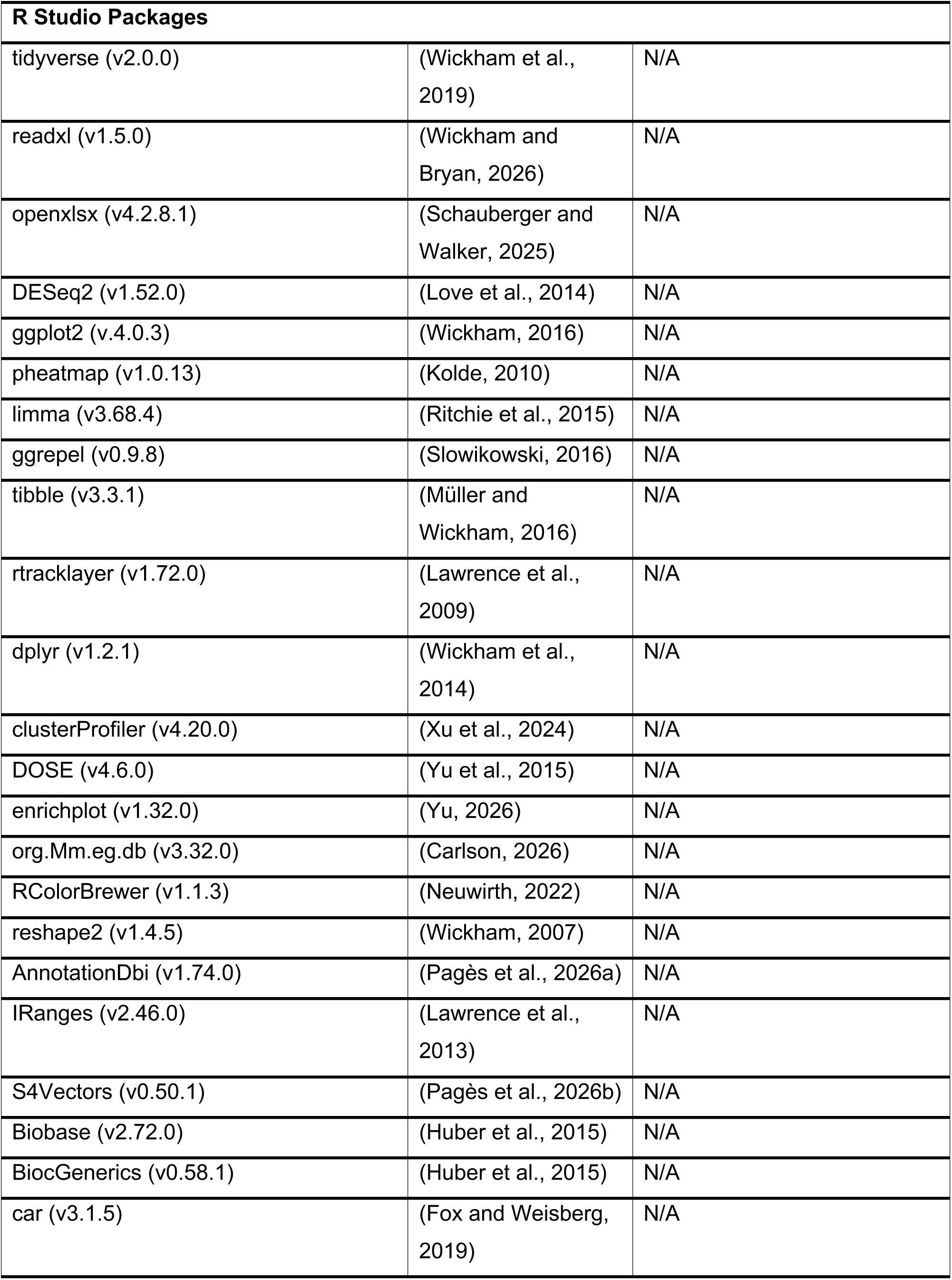
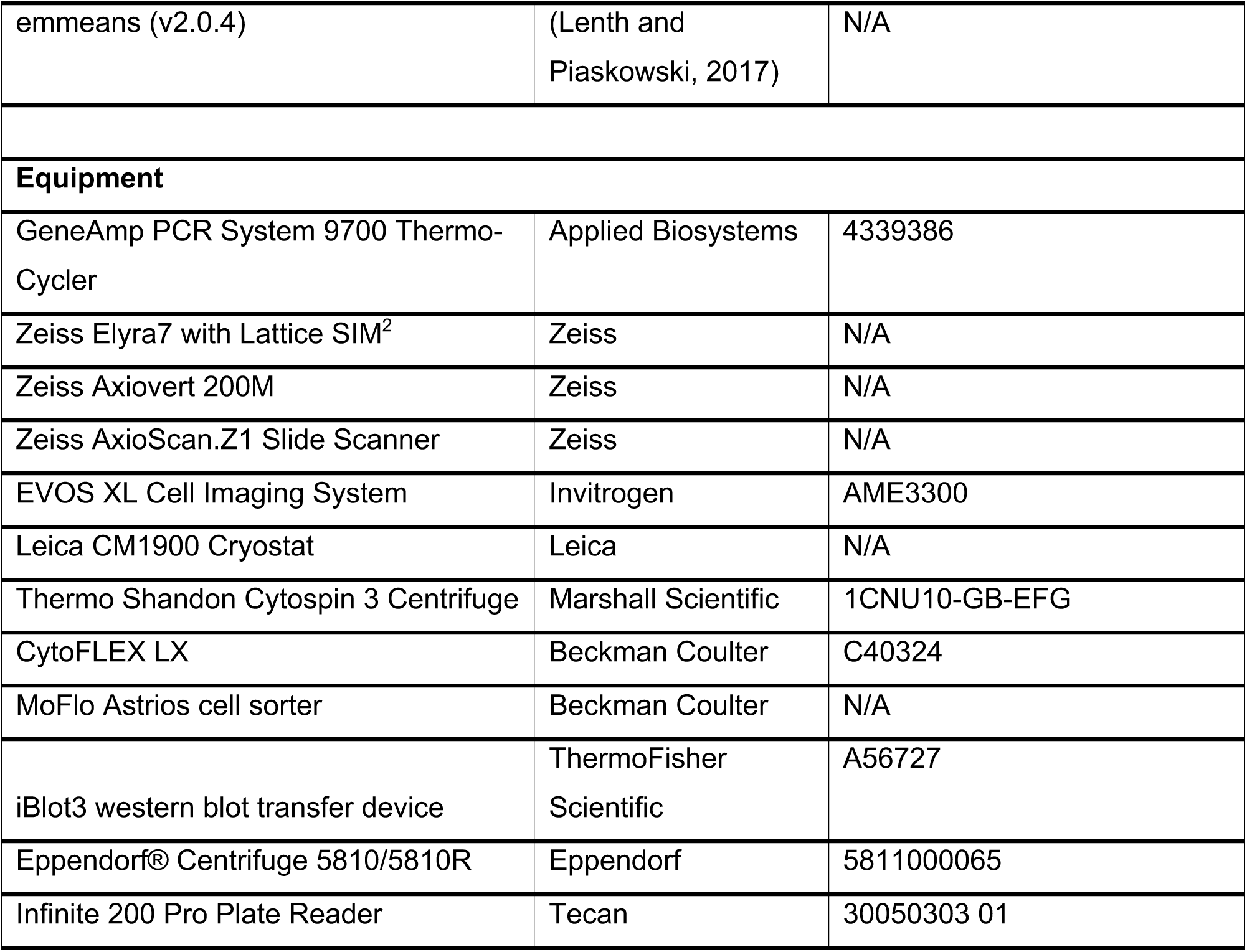

### Mice and genotyping

All animal work was carried out with ethical approval from the University of York under UK Home Office approved Project Licence PP2435190 where the animals were maintained under specific pathogen free conditions. CIZ1-null mice were generated from *Ciz1*+/- C57BL/6 embryonic stem cell clone IST13830B6 (Texas A&M Institute for Genomic Medicine) harboring a neomycin resistance gene trap insertion into intron 1 (Ridings-Figueroa et al., 2017), and maintained by interbreeding over multiple generations.

CIZ1 genotypes were confirmed by PCR following DNA extraction using the PCRBio Rapid Extract Lysis Kit (PCR Biosystems) from ear notches. Briefly, tissue samples were mixed with 10 µl 5x PCRBio Rapid Extract buffer A, 5 µl 10x PCRBio Rapid Extract buffer B, and 35 µl nuclease-free water (total 50 µl per tissue sample). Samples were lysed at 75°C for 45 minutes with vortexing every 15 minutes, then placed at 95°C for 10 minutes to deactivate the protease. Samples were centrifuged briefly to remove condensation from the lid, and 50 µl TE buffer (Invitrogen) was added per sample, and centrifuged at 10,000 x g for 1 minute to pellet the residual tissue. The supernatant was transferred to a clean Eppendorf and stored at 4°C. For PCR, 10 µl 2x KAPA2G Fast ReadyMix (Roche), 1 µl 10 µM IVS1F primer (GTGGCGTTGGCTATATCTGC), 1 µl 10 µM IVS1R primer (GTTGAACATGGTGGCTGAAG), and 1 µl 10 µM VR primer (CCAATAAACCCTCTTGCAGTTGC), 5 µl nuclease-free water (Invitrogen), and 2 µl extracted DNA were mixed. Positive DNA and a negative water control were included. PCR reactions were placed in a GeneAmp PCR System 9700 Thermo-Cycler (Applied Biosystems) under the following PCR cycle: Melting hold (95°C, 3 minutes); 29 cycles of PCR amplification (melting: 95°C, 15 seconds; annealing: 60°C, 15 seconds; elongation: 72°C, 5 seconds), final elongation (72°C, 1 minute) and a final hold (4°C, 10 minutes). 10µl PCR products alongside 5 µl ØX174 DNA-HaeIII Digest ladder (New England Biolabs) were separated and analysed on a 1.5% w/v agarose (Melford) gel made with 1x TBE (89 mM Tris, 89 mM Boric acid, 2 mM EDTA) supplemented with a 1:10,000 dilution of SYBR™ Safe DNA Gel Stain (Invitrogen). Electrophoresis was carried out in a gel tank containing 1x TBE at 90 V, for 1 hour.

### Splenocyte isolation

Spleens were collected within 5 minutes of euthanasia into ice-cold PBS, weighed, then mashed through a 70 µm^2^ cell strainer with 15 ml ice-cold RPMI 1640/1% Penicillin-Streptomycin/10% FBS media. The splenocyte suspension was pelleted at 300 x g, 4°C for 5 minutes, and the supernatant discarded. 1 ml red blood cell lysis buffer (Sigma-Aldrich) was added and the pellet resuspended gently, then left on ice for 5 minutes, before dilution to 10 ml with PBS, and pelleted to discard the supernatant. Pellets were resuspended in 10 ml media. 100 µl aliquots were reserved for counting using a haemocytometer, after 1:10 dilution in PBS and 1:1 mix with Trypan blue.

### MACS sorting

After splenocyte isolation and red blood cell lysis, negative B cell isolation was achieved using the Pan B Cell Isolation Kit II by Miltenyi Biotec with MACS LS separation columns, following the protocol supplied. From an input of 10^7^ splenocytes, ∼3–5 x 10^6^ B cells were isolated. Cell viability was quantified using a haemocytometer after staining 1:1 with Trypan blue, and B cell enrichment by immunostaining cytospun samples with α-B220.

### *In vitro* splenocyte activation

10^6^ splenocytes were plated per well of a flat-bottomed 96-well plate with 100 µl RPMI 1640/10% heat-inactivated FBS/1% Penicillin-Streptomycin/2mM L-glutamine media. 100 µl media supplemented with 2 µg/ml LPS (Sigma-Aldrich) or αCD3 (BioLegend) was added per well, giving a final working concentration of 1 µg/ml. Splenocytes were incubated at 37°C, 5% CO_2_ for 24 and 48 hours, with daily imaging on an EVOS (serial number: L0809-1493-043, software 9978) fitted with a 20X AGM Plan LWD PH (AMEP-4634) objective.

### Bone marrow cell isolation

Tibiae, femurs, ilia bones and spines were dissected from individual mice at 8–10 weeks, cleaned of loose tissue and spinal cord, and collected in ice-cold PBS/2% FBS. Bones were crushed with a pestle and mortar with 30 ml ice-cold PBS/2% FBS and strained through a 70 µm^2^ cell strainer. The cell suspension was pelleted at 300 x g, 4°C for 5 minutes, and the supernatant discarded. To lyse red blood cells, 3 ml PBS and 5 ml ammonium chloride solution (STEMCELL Technologies) was added without disturbing the pellet and incubated on ice for 5 minutes. Pellets were then resuspended by gentle mixing and incubated for a further 5 minutes on ice. Samples were pelleted at 300 x g, 4°C for 5 minutes and the supernatant discarded.

Pellets were resuspended in 10 ml PBS/2% FBS and split into fractions, then pelleted at 300 x g, 4°C for 5 minutes and the supernatant discarded before proceeding with antibody staining. Fractions were 40% ESLAM panel (for LT-HSCs), 40% MPP panel (ST-HSCs and MPP-Lys), and 20% for single stain controls. Due to their rarity, if ESLAM LT-HSCs were to be sorted, bone marrow cell suspensions were enriched for stem and progenitor cells first. Following red blood cell lysis, bone marrow pellets were resuspended in 500 µl 2% FBS/PBS. Enrichment was achieved using the EasySep Mouse Hematopoietic Progenitor Cell Isolation Kit (STEMCELL Technologies) with MACS magnets. 10 µl EasySep HSPC Isolation Cocktail was mixed to each sample and incubated on ice for 15 minutes. Then, 15 µl of thoroughly vortexed EasySep RapidSpheres was mixed to each sample and incubated on ice for another 15 minutes.

Samples were topped up with 2 ml 2% FBS/PBS, mixed gently, then placed into a MACS magnet at room temperature for 3 minutes. The enriched cell suspension was collected by inverting the magnet and tubes together. An additional 2 ml 2% FBS/PBS was added to the sample tube containing the beads to provide an additional wash ending up with ∼4 ml enriched cell suspension. Tubes containing the magnetic beads and lineage cells were discarded. Enriched cells were pelleted and the supernatant discarded.

### Cell staining and flow cytometry

*<u>Splenocytes:</u>* For each antibody panel, 5 x 10^6^ splenocytes were pelleted at 300 x g, 4°C for 5 minutes, and the supernatant discarded. For identification of dead cells, pellets were resuspended in 100 µl Zombie Aqua (1:1000) in PBS and placed on ice for 10 minutes, protected from light. Cells were washed with 1 ml RPMI 1640/1% Penicillin-Streptomycin/10% FBS, pelleted, and resuspended in 100 µl T and B cell antibody cocktail (CD4 PerCP-Cy5.5, CD8 Pacific Blue, TCRβ PE-Cy7, CD19 APC-Cy7, B220 APC) and left on ice for 30 minutes, protected from light. Cells were washed with 1 ml PBS/2% FBS, pelleted and the supernatant discarded. Pellets were resuspended in 200 µl PBS/2% FBS and plated onto a U-bottomed 96-well plate kept on ice.

*<u>Bone marrow:</u>* Pellets were resuspended in 100 µl of either primary ESLAM (for LT-HSCs) antibody cocktail (CD45 FITC, EPCR PE, CD150 PE-Cy7, Sca-1 BV605, CD48 APC, CD44 Biotin, CD5 Biotin, CD19 Biotin, TER119 Biotin, CD45R/B220 Biotin) or primary MPP (for ST-HSCs and MPP-Lys) antibody cocktail (c-Kit APC-Cy7, Sca-1 BV605, CD135 BV421, CD48 APC, CD150 PE-Cy7, CD44 Biotin, CD5 Biotin, CD19 Biotin, TER119 Biotin, CD45R/B220 Biotin) and incubated on ice for 30 minutes protected from light. 1 ml PBS/2% FBS was added to each sample, then pelleted and the supernatant discarded. Pellets were resuspended in 100 µl of either secondary ESLAM antibody cocktail (Streptavidin BV510) or secondary MPP antibody cocktail (Streptavidin BV785) and incubated on ice for 30 minutes, protected from light. 1 ml PBS/2% FBS was added to each sample, then pelleted and the supernatant discarded. All fractions were thoroughly resuspended in 500 µl 1:1000 7AAD for 5 minutes, then filtered through a blue-cap 5-ml polystyrene FACS tube and plated in a U-bottomed 96-well plate on ice, protected from light. Cell types were sorted on the basis of the following markers, LT-HSCs: Lineage^−^ CD48^−^ CD150^+^ CD45^+^ EPCR^+^ (Kent et al., 2009), ST-HSCs: Lineage^−^ c-Kit^+^ Sca-1^+^ CD135^−^ CD48^−^ CD150^−^, and MPP-Ly: Lineage^−^ c-Kit^+^ Sca-1^+^ CD135^+^. Alongside, single stain controls were prepared either by mixing 1.5 ml PBS/2% FBS with 4 drops of UltraComp Ebeads (Invitrogen) or by using reserved live cells split into 100 µl fractions. One fraction remained unstained while the rest received 0.5 µl of individual antibody then placed on ice for 30 minutes, protected from light. For Zombie Aqua or 7AAD live/dead single stain controls, reserved cells were always used and placed on a 65°C heat block for 10 minutes to induce cell death prior to antibody addition. Single stains were plated on the sample plate and kept on ice, ready for cytometric detection. Plates were analysed on a Beckman Coulter CytoFLEX LX with 6 lasers (355nm to 808nm) within 3–4 hours of isolation, calibrated using single stain controls. Cells were gated on live, single cells before proceeding to antibody gating strategies.

### Spleen cryosections

Whole spleens were submerged in 4% PFA and fixed at 4°C for 5 hours, rinsed in PBS thoroughly and submerged in 30% (w/v) sucrose/PBS solution at 4°C overnight. Spleens were cut in half transversally, embedded in OCT embedding matrix (CellPath) and flash frozen over a dry ice slurry made with ethanol. Embedded spleens were cut on a Leica CM 1900 cryostat at – 20°C as 5 µm cryosections onto SuperFrost microscope slides (Epredia). Sections were left to air dry for 20 minutes before staining or storing at –80°C. For staining, hydrophobic rings were drawn around the sections using a peroxidase-anti-peroxidase (PAP) pen (Merck) and rehydrated in BSA antibody buffer (1% BSA, 0.1% Triton-X-100, 2% SDS) 3 times for 5 minutes each. Sections were then permeabilised with 2% Triton-X in PBS for 30 minutes at room temperature, then washed 3 times in BSA antibody buffer for 5 minutes each before incubation with antibody as described below.

### Immunofluorescence staining

Approximately 100,000 cells were fixed in suspension by adding them to 200 µl 4% PFA for 10 minutes at room temperature, and the sample loaded on a Shandon Cytospinner 3 centrifuge, and spun onto SuperFrost microscope slides (Epredia) at 20 x g for 5 minutes. Slides were air dried and a hydrophobic ring drawn around the cells with a PAP pen. Slides were washed 3 times with PBS for 2 minutes each, and then with CSK buffer (10 mM PIPES pH6.8, 100 mM NaCl, 1 mM MgCl_2_, 300 mM sucrose, 1 mM EGTA) supplemented with triton X-100 (final concentration 0.1%) for 1 minute, then again with PBS. Samples were blocked in BSA antibody buffer for 30 minutes before staining. In a humidified chamber, 20 µl primary antibody cocktail prepared in BSA antibody buffer was added to cell samples, the chamber sealed and incubated at 37°C for 1.5 hours. Samples were washed in BSA antibody buffer 3 times, then incubated with secondary antibody cocktail prepared in BSA antibody buffer, at 37°C for 1 hour protected from light. Samples were washed sequentially in BSA antibody buffer, PBS and HPLC water (Fisher Scientific) then mounted with DAPI containing Vectashield (Vector labs) and covered with a glass coverslip.

### Fluorescence microscopy

Stained cytospun samples were imaged on a Zeiss Axiovert 200M microscope fitted with a 63x/1.40 Plan Apochromat objective and Zeiss filter sets 1, 10, and 15 (BP365/12 FT 395 LP397, BP450–490 FT510 BP515–565, BP546/12 FT580 LP590) using Axiocam 506 mono and ZEN Blue 3.1 acquisition software. For super-resolution microscopy, images were taken as Z-stacks on a Carl Zeiss Elyra 7 super-resolution microscope fitted with an alpha Plan-Apochromat 63x/1.46 Oil Korr M27-Oil objective and laser sets 405, 488, and 561 nm, using ZEN blue 3.13 acquisition and processing software. All channels for each image underwent SIM^2^ processing. Zeiss Immersol 518F was used for all cellular imaging. Consistent exposure time parameters were used for each antibody within an experiment to generate image sets with meaningful relative intensity, unless stated otherwise. Internally controlled image sets were quantified using ImageJ2 software (version 2.16.0/1.54s) performed on raw unmodified images. For reproduction purposes, some images were adjusted using ImageJ2 software, in all cases to preserve quantified relationships. Stained cryosections were imaged using an Axio Scan.Z1 slide scanner with a Plan-Apochromat 40x/0.95 Korr M27 objective. Due to differences in fluorescent intensity across different spleen isolates, images were not identically captured. No fluorescent intensity quantifications have been made on fluorescent cryosections imaged by Slidescanner.

### ELISA

Blood samples (∼200 µl) were collected in EDTA-tubes from mice following euthanasia. Blood was spun at 1800 x g for 8 minutes, and the supernatant (plasma) collected and stored at – 20°C. Plasma was diluted 1:5000 with 100 mM BupH Carbonate-Bicarbonate coating buffer, and 200 µl added into row A of a 96-well MaxiSorp plate (Invitrogen). Rows B–H were filled with 100 µl coating buffer. 100 µl of each sample was serially diluted 2-fold by transfer into a recipient well containing 100 µl of coating buffer. Plates were sealed and left at 4°C for 2 hours to allow binding. Wells were washed 4 times with 200 µl PBS/0.2% Tween-20 wash buffer, then blocked with 100 µl PBS/1% BSA (Sigma)/0.3% Tween-20 blocking buffer at 37°C for 1 hour, sealed. Wells were replaced with 100 µl 1:10,000 α-mouse IgG-HRP antibody prepared in blocking buffer and incubated at 37°C for 1 hour. Wells were washed 4 times then developed using 100 µl room temperature TMB substrate (SeraCare) for 5 minutes, followed by 100 µl STOP solution and then read at 450 nm on an Infinite 200 Pro Plate Reader within 30 minutes of development. All plates included the same positive control sample to create a standardisation curve and a negative control blank well.

### Whole cell lysates and western blots

10^6^ cells were spun down at 300 x g for 5 minutes, and the supernatant discarded. Pellets were resuspended in 1x SDS-PAGE denaturing buffer (4% SDS, 15% glycerol, 1.7% β-mercaptoethanol, 75 mM Tris pH6.8, pinch of bromoethanol blue) and heated at 95°C for 10 minutes, with repeated vortexing in between to thoroughly lyse and denature the samples. Samples were then frozen at –80°C or separated by electrophoresis through a 4–15% gradient gel (BioRad). Gels were transferred onto a nitrocellulose membrane using the iBlot3 system (Invitrogen). Blots were blocked with 5% non-fat dried milk powder (Apollo Scientific) in PBS with 0.1% Tween-20, then incubated with primary antibodies either overnight at 4°C or at room temperature for 2 hours with gentle agitation. Blots were washed and probed with HRP-conjugated anti-species secondary antibodies (Jackson ImmunoResearch) for 1 hour at room temperature and imaged using Westar Antares ECL substrate (Cyanagen) with a Syngene PXi chemiluminescence imaging system.

### RNA isolation

Young naive and LPS-activated splenocytes (4×10^6^ cells) or enriched naive B cells (2×10^6^ cells) were pelleted at 300 x g for 5 minutes, supernatant discarded, and 1 ml Trizol (Invitrogen) added to the cell pellets. Adult spleens from 15+ month old animals were frozen, and shavings were placed directly into 1 ml Trizol (Invitrogen). Trizol samples were mixed then frozen at – 80°C, or processed immediately to extract RNA as recommended by the supplier. Unless otherwise stated, all processing was done at room temperature. Briefly, after adding 0.2 ml chloroform, the samples were mixed thoroughly by hand for 15 seconds, incubated for 3 minutes, then centrifuged at 12,000 x g for 15 minutes. 450 µl of the upper aqueous phase was carefully transferred to a fresh tube, an equivalent volume of isopropyl alcohol added, mixed, incubated for 10 minutes then centrifuged at 12,000 x g for 10 minutes. The RNA pellet was washed with 75% ethanol in DEPC water, dried briefly and resuspended in 20–50 µl DEPC water (Ambion) by heating to 55°C for 10 minutes before gently pipetting. Samples were then kept on ice, concentration and quality tested using a NanoDrop Spectrophotometer (ND-1000) and stored at –80°C.

### RNA-sequencing and expression analysis

RNA samples were taken through Zymo Research RNA Clean and Concentrator columns with an on-column DNase digestion and then RNA quality assessed on an Agilent Technologies Bioanalyzer 2100. Libraries were prepared for sequencing using the NEBNext® Ultra II Directional RNA Library prep kit for Illumina in conjunction with the NEBNext® Poly(A) mRNA Magnetic Isolation Module (New England Biolabs), according to the manufacturer’s instructions. Amplification of libraries used NEBNext® Multiplex Oligos for Illumina Unique Dual Index Primer Pairs (New England Biolabs). Libraries were pooled at equimolar ratios and sequenced using paired-end 150 base sequencing at Azenta Life Sciences, using an Illumina NovaSeq X to generate 20 million reads per sample. The Viking cluster (University of York) was used to perform further analysis. Output FastQ files were quality-checked using FastQC v0.12.1-Java-11 (Andrews, 2010) pre- and post-trimming. Cutadapt v4.2-GCCcore-11.3.0 (Martin, 2011) was used to remove sequences with a Phred score <20 (less than 99% confidence) and smaller than 75bp. Sequences were mapped and aligned at the gene level using STAR v2.7.10b-GCC-11.3.0 (Dobin et al., 2013), with the “--quantMode” flag, against mouse primary assembly genome GRCm39 alongside GRCm39 vM36 primary annotation GTF file (Mudge et al., 2025) providing a “ReadsPerGene.out.tab” count table file for each sample. Genes which expressed a sum count <10 across all replicate samples, were converted to 0 to mitigate false-positive significant differential expression scores when absolute expression levels are low, and potentially inconsistent across samples. Count table differential expression analysis was achieved using DESeq2 v1.49.4 (Love et al., 2014) in R studio v4.5.0 (PositTeam, 2025). The naive and activated female splenocyte data was collected as two independent experiments separated by over five years (two sets of n=2 for each genotype and condition). During differential expression analysis of naive and activated female splenocytes, batch was included as a covariate in the design formula (∼batch + condition). This allowed differential expression between conditions to be estimated while controlling for variation attributable to sequencing batch. For all other experiments sample preparation and sequencing was performed together. GRCm39 vM36 primary annotation GTF file (Mudge et al., 2025) was used for sequence mapping and to annotate the DESeq2 differential expression output to include Gene ID, Gene Name, Chromosome, Start position, End position, RNA type, Strand, Base Mean, Log_2_ Fold Change (Log_2_FC), *p*-value, and *q*-value.

### Statistics and Data visualization

Significant DEGs were determined by *p-* or *q-*value thresholds, as indicated. To visualise location of affected genes, data was split by chromosomes, and log_2_FC (y-axis) was plotted against gene start position (x-axis). Data was analysed and plotted in Microsoft Excel (version 16.107.3). Confirmatory Volcano and PCA plots were generated using R Studio and Stats v4.5.0, DESeq2 v1.49.4, and ggplot2 v3.5.2. packages. Other graphical illustrations were either generated using Excel or in R studio v4.5.0 (PositTeam, 2025). Box and whisker plots were generated in Excel and show inter-quartile range with outliers, means represented by ‘X’, and the median by a line. For heat maps, log2 fold changes and Z-scores were calculated from TPM replicate means, by calculating average and standard deviation across a comparison set, for each gene.

For comparison between two datasets, a two-tailed Student’s T-test accounting for equal or unequal variance was performed in Excel. To analyse the effects of sex and genotype on spleen weight, IgG plasma levels, and gene transcript levels, data was analysed using a Two-Way Analysis of Variance (ANOVA). Due to unequal sample sizes across experimental groups, a Type III Sum of Squares error structure was utilised to prevent bias from unbalanced data.

Significant interactions were further evaluated using a post-hoc pairwise comparison with Estimated Marginal Means (EMMeans), applying Tukey’s Honestly Significant Difference adjustment to correct for multiple comparisons. Alpha levels for all primary and post-hoc analyses were set beforehand at 0.05. Statistical analyses were performed in R studio v4.5.0 (PositTeam, 2025) utilising the “car” and “emmeans” packages.

Statistical tests used in each analysis are stated in the figure legend with *p*-values, where asterisks indicate \**p* ≤ 0.05, \*\**p* ≤ 0.01, \*\*\**p* ≤ 0.001.

### DEG excess enrichment (EE)-score and cluster plot analysis

To calculate DEG EE-scores each chromosome was divided into 6 Mb bins, overlapping by 2 Mb that were sequentially numbered 0, 2, 4, etc so that the bin number coincides with its start position in Mb. For each chromosome, the following parameters were calculated: 1) the total number of DEGs (sum DEGs); 2) the total number of genes (sum genes); 3) the total minimum number of bins required to span the length of the chromosome (total bins). These were used to calculate the expected DEG frequency (sum DEGs ÷ total bins) and the expected gene frequency (sum genes ÷ total bins) for each chromosome. For each bin on each chromosome the actual number of DEGs and genes were used to calculate their DEG and gene enrichment scores (E-Scores), where E-Score = observed ÷ expected frequency. To understand which bins are more enriched for DEGs than genes, we calculated the DEG EE-Score which is the difference in DEG E-Score over gene E-Score. A DEG EE-Score ≥ 3 was used to select for bins with high enrichment for DEGs. Clusters between the sexes are considered overlapping if one or more bins that define a cluster covers common range with the range of another bin in the opposite sex. For example: if a female cluster spans bins 0 and 2 (range: 0–8 Mb) and a male cluster spans bins 4 and 6 (range: 4–12 Mb) on the same chromosome, they would be considered overlapping as they span a common area (4–8 Mb), even though they do not share the same bin number.

### GSEA

Gene Set Enrichment Analysis (GSEA) was performed using the R/Bioconductor packages org.Mm.eg.db for genome wide annotation for mouse (Carlson, 2026), ClusterProfiler v4.20.0 (Xu et al., 2024) and DOSE v4.6.0 (Yu et al., 2015) against mouse biological processes (BP) gene sets using cut-off *p*-value<0.05 and by using a fixed random seed “123” to ensure reproducibility in R Studio v4.5.0 (PositTeam, 2025). Alternatively, GSEA was performed using the online GSEA-MSigDB tool (version mouse MSigDB v2026.1.Mm) against mouse M5: Ontology gene sets (FDR *q*-value<0.05) (Subramanian et al., 2005).

## Supplemental Tables

Supplemental Table 1 – Spleen (mg) and body (g) weights, and derived SWI.

## Supplemental Datasets

Supplemental dataset 1 – CIZ1null vs WT Adult female spleen 15m

- Tab 1 – Count table + TPMs

- Tab 2 – GSEA

- Tab 3 – DESeq2 analysis

- Tab 4–23 – Cluster analysis with biotype proportions per chromosome

- Tab 24 – Cluster boundaries

Supplemental dataset 2 – WT LPS vs NAIVE 8–10wk female splenocytes

- Tab 1 – Count table + TPMs

- Tab 2 – Transcription factor gene sig

- Tab 3 – DESeq2 analysis

- Tab 4–23 – Cluster analysis with biotype proportions per chromosome

Supplemental dataset 3 –CIZ1null LPS vs NAIVE 8–10wk female splenocytes

- Tab 1 – Count table + TPMs

- Tab 2 – Transcription factor gene sig

- Tab 3 – Missing 2326 DEGs

- Tab 4 – DESeq2 analysis

- Tab 5–24 – Cluster analysis with biotype proportions per chromosome

Supplemental dataset 4 – NAIVE CIZ1null vs WT 8-10wk female splenocytes

- Tab 1 – Count table + TPMs

- Tab 2 – GSEA

- Tab 3 – DESeq2 analysis

- Tab 4–23 – Cluster analysis with biotype proportions per chromosome

Supplemental dataset 5 – LPS CIZ1null vs WT 8-10wk female splenocytes

- Tab 1 – Count table + TPMs

- Tab 2 – GSEA

- Tab 3 – DESeq2 analysis

- Tab 4–23 – Cluster analysis with biotype proportions per chromosome

Supplemental dataset 6 – CIZ1null vs WT 8-10wk female naive B cells

- Tab 1 – Count table + TPMs

- Tab 2 – GSEA

- Tab 3 – Transcription factor gene sig

- Tab 4 – DESeq2 analysis

- Tab 5–24 – Cluster analysis with biotype proportions per chromosome

- Tab 25 – Escape gene cluster analysis

- Tab 26 – Cluster boundaries

Supplemental dataset 7 – CIZ1null vs WT 8-10wk male naive B cells

- Tab 1 – Count table + TPMs

- Tab 2 – GSEA

- Tab 3 – Transcription factor gene sig

- Tab 4 – DESeq2 analysis

- Tab 5–25 – Cluster analysis with biotype proportions per chromosome

- Tab 26 – Escape gene cluster analysis

- Tab 27 – Cluster boundaries

Supplemental dataset 8 – Cluster map of CIZ1null vs WT 8-10wk female and male naive B cells

- Tab 1 – Cluster map

---

**S.Fig 1.**
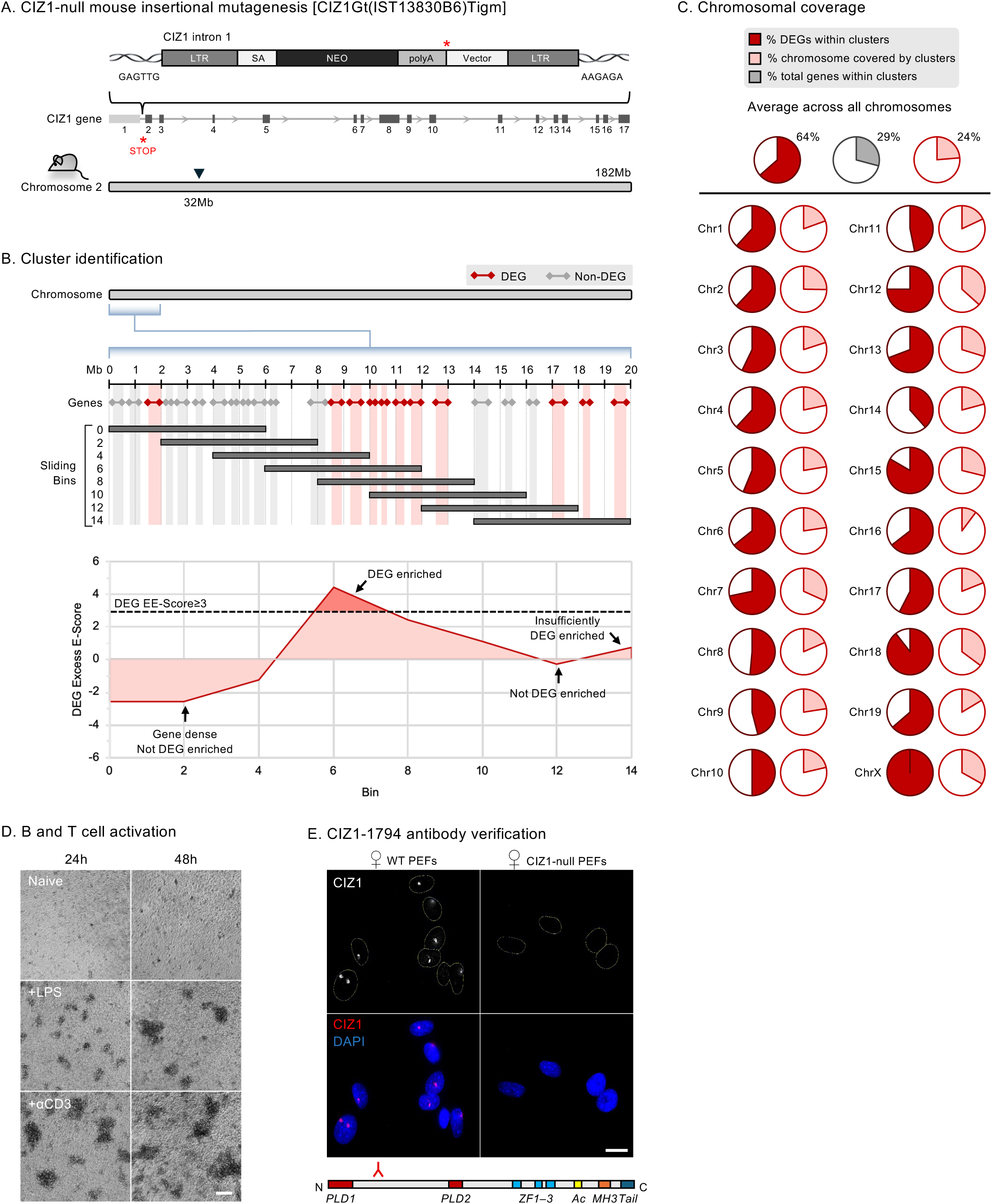
(Supplemental to Figs 1 and 2) A) Insertional mutagenesis strategy at CIZ1 intron 1 to generate the CIZ1-null mouse line. B) Schematic of the sliding bin strategy to identify DEG-enriched bins. Bins with DEG EE–Score≥3 were included in the analysis. C) Pie charts of 15+ month old female CIZ1-null vs WT spleens showing proportion of genes that are differentially expressed within cluster sites, the proportion of chromosomes that are covered by clusters (EE-Score≥3), and the overall proportion of genes found within clusters (EE-Score≥3) . D) Brightfield images of female WT total splenocytes following either LPS or αCD3 activation for 24 or 48 hours. Scale bar is 100 µm. E) Validation of CIZ1-1794 antibody to recognise CIZ1 at the Xi (grey/red) in female WT primary embryonic fibroblasts (PEFs). Female CIZ1-null PEFs show no signal. Scale bar is 20 µm. Below, schematic of CIZ1 protein domains (Byrom et al., 2026), showing region recognised by anti-CIZ1 rabbit polyclonal antibody 1794.

**S.Fig 2.**
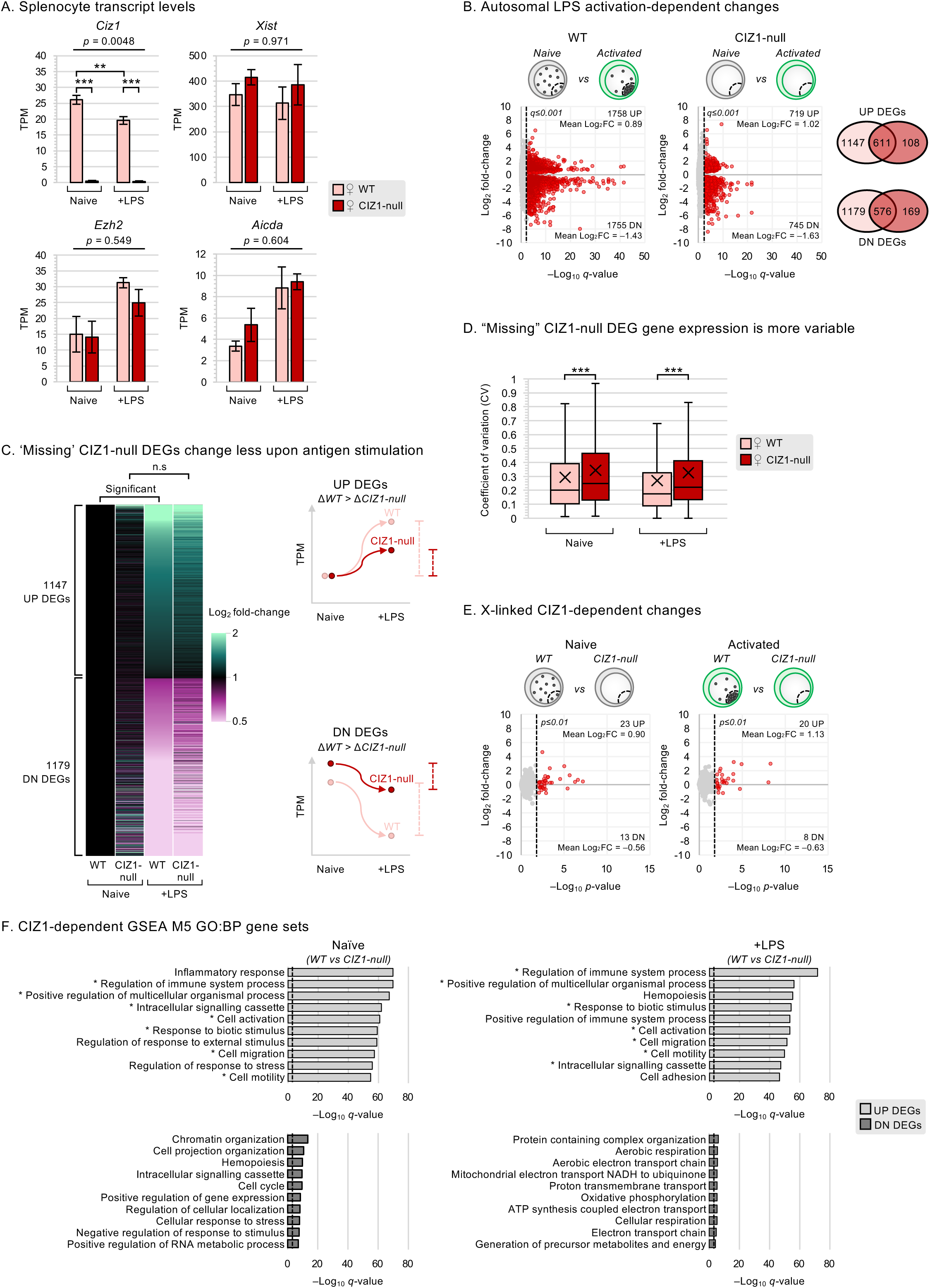
(Supplemental to Fig 3). A) Transcript levels of *Ciz1, Xist, Ezh2, and* AID *(Aicda)* in female WT and CIZ1-null, naive and LPS-activated splenocytes of 8–10 week old mice. Analysis by Two-Way ANOVA. B) Volcano plots showing activation-dependent changes in autosomal genes (*q*≤0.001) in female splenocytes from 8–10 week old mice, showing WT (left) and CIZ1-null (right). Venn diagrams show mutual DEGs between WT (pink) and CIZ1-null (red) UP and DN DEG populations. C) Heatmap showing TPM log_2_ fold-change for the ‘missing’ 2326 genes in female WT and CIZ1-null, naive and LPS-activated splenocytes. Average TPMs across four replicates were normalised to female WT naive splenocytes. Right, models of DEG trajectories for WT and CIZ1-null, UP and DN DEGs. D) Coefficients of variation of the ’missing’ 2326 genes (calculated as the standard deviation divided by the mean) for female WT (pink) and CIZ1-null (red), naive and activated splenocytes. Statistical analysis performed by Student’s T-test. E) Volcano plots showing X-linked CIZ1-dependent changes (p≤0.01) in female splenocytes from 8–10 week old mice, showing naive (left) and activated (right). F) The top 10 UP and DN gene expression pathways (GSEA M5 Biological Processes) affected by the absence of CIZ1 in naive and LPS-activated splenocytes (autosomal and X-linked DEGs combined). Dotted lines depict significance threshold set to *q*≤0.001.

**S.Fig 3.**
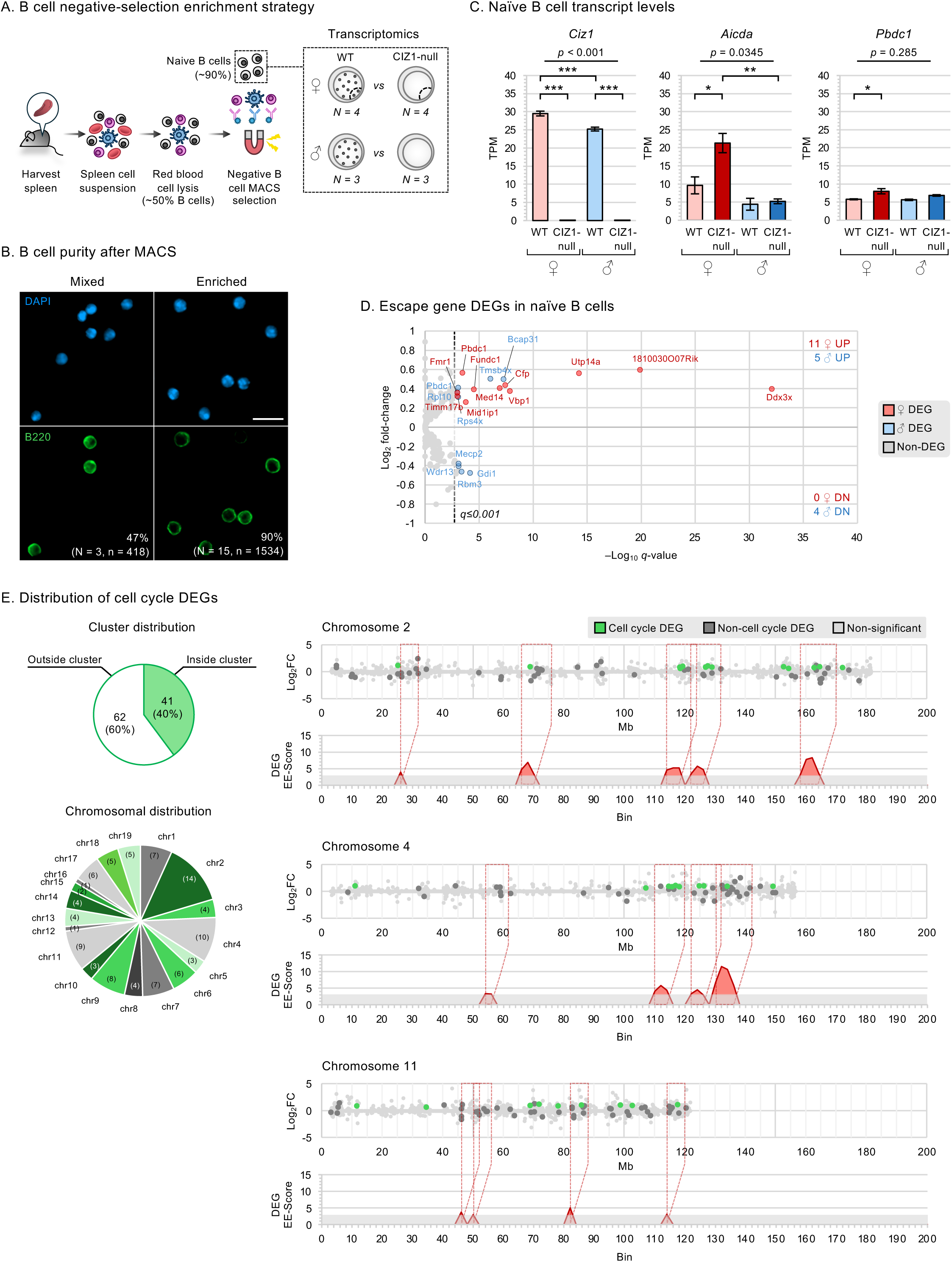
(Supplemental to Fig 4 and analysis of cell cycle genes) A) B cell purification strategy by negative-selection, and outline of B cell transcriptomes generated. N represents number of mice. B) Example images illustrating B cell purity (%) pre- and post-MACS sorting, scored based on immunofluorescence signal of pan-B cell marker B220 (green) and DAPI (blue). N = number of animals, n = number of cells. Scale bar is 10 µm. C) Transcript levels of *Ciz1*, AID (*Aicda*) and *Pbdc1* in WT and CIZ1-null, female and male naive B cells. Analysis by Two-Way ANOVA. D) Volcano plot showing escape genes in naive B cells, highlighting significant (q≤0.001) CIZ1-dependent escape DEGs in females (red) and males (blue). E) Cluster localization of the 103 cell cycle-related differentially expressed DEGs derived from the top five UP regulated gene sets in GSEA M5 biological processes returned by female naive B cells (Fig.5D). Below, their chromosomal origin with number of cell-cycling genes identified in parentheses. Right, locations of the top three female naive B cell chromosomes (2, 4 and 11) enriched for cell cycle DEGs (green). Below, EE-Score≥3 derived gene cluster sites.

**S.Fig 4.**
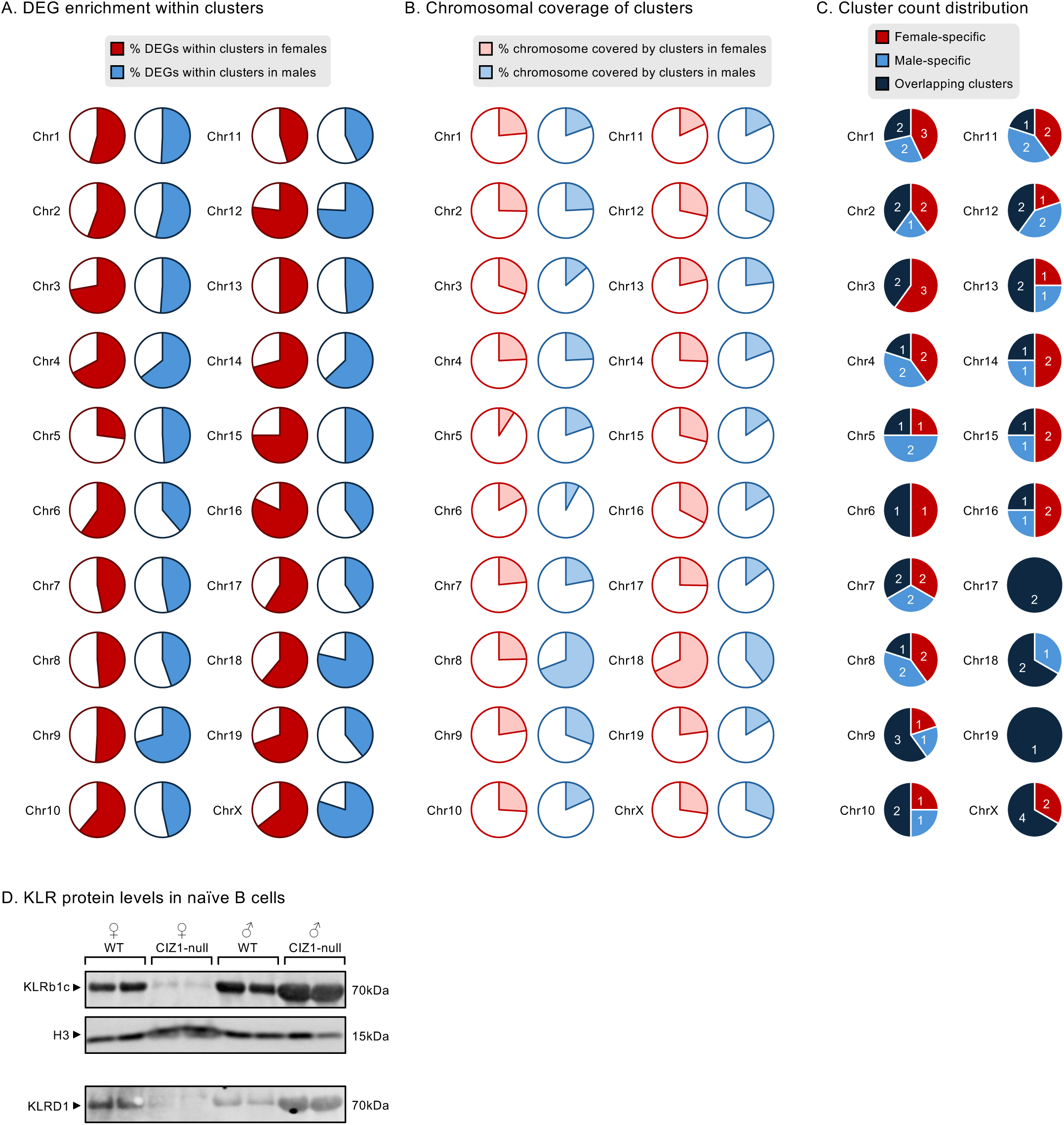
(Supplemental to Figs 5 and 6) A) Percentage of CIZ1-dependent DEGs found within clusters in individual chromosomes in females (red) and males (blue). B) Percentage of chromosome covered by CIZ1-dependent DEG-defined clusters in females (light red) and males (light blue). C) Distribution of CIZ1-dependent DEG-defined clusters across all chromosomes in female and male naive B cells. Clusters are defined as female-specific (red), male-specific (light blue) or overlapping between the sexes (dark blue). D) Western blots of naive B cell whole cell lysates showing KLRb1c alongside histone H3 loading control. KLRD1 protein was blotted on a separate membrane.

**Supplemental Table 1.**
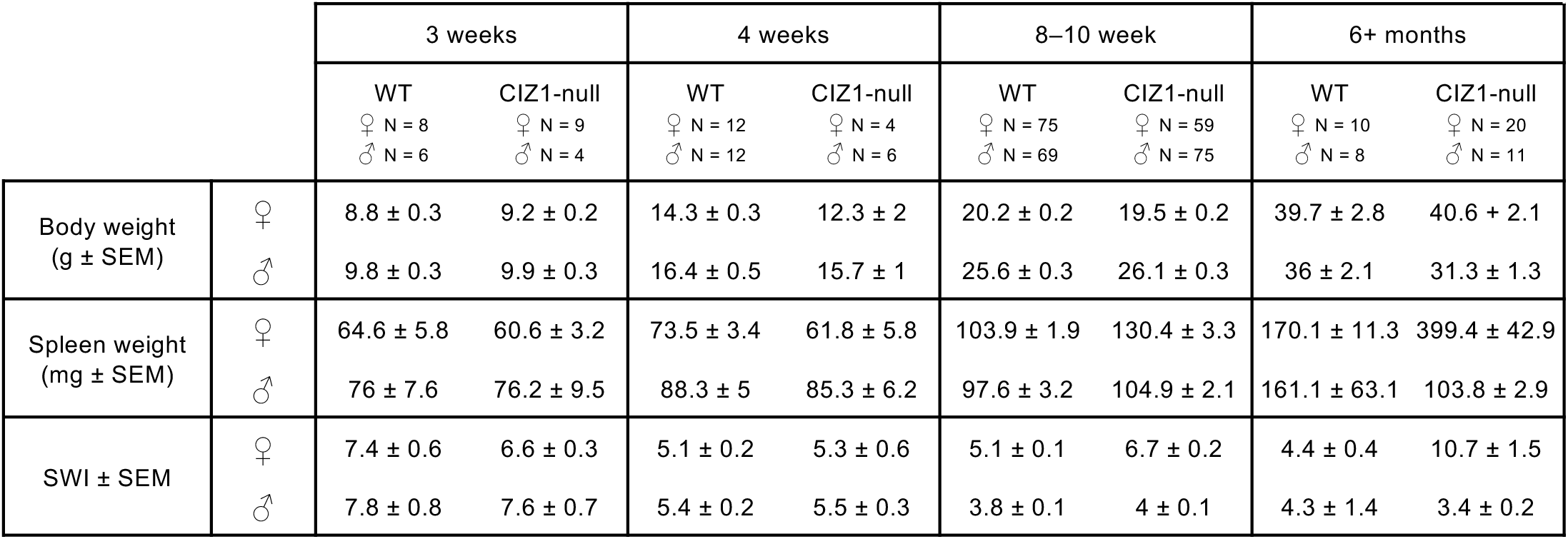
Body, spleen weights and spleen weight index (SWI) of mice used at 3, 4, 8–10 weeks and 6+ months (±SEM). Animals aged 6+ months were ex-breeders. Number of animals used (N) are indicated.

